# Low grade inflammation in the epileptic hippocampus contrasts with explosive inflammation occurring in the acute phase following *status epilepticus* in rats: translation to patients with epilepsy

**DOI:** 10.1101/2021.03.25.436701

**Authors:** Nadia Gasmi, Fabrice P. Navarro, Michaël Ogier, Amor Belmeguenaï, Thomas Lieutaud, Béatrice Georges, Jacques Bodennec, Marc Guénot, Nathalie Streichenberger, Philippe Ryvlin, Sylvain Rheims, Laurent Bezin

## Abstract

There is still a lack of robust data, acquired identically and reliably from tissues either surgically resected from patients with mesial temporal lobe epilepsy (mTLE) or collected in animal models, to answer the question of whether the degree of inflammation of the hippocampus differs between mTLE patients, and between epilepsy and epileptogenesis. Here, using highly calibrated RTqPCR, we show that neuroinflammatory marker expression was highly variable in the hippocampus and the amygdala of mTLE patients. This variability was not associated with gender, age, duration of epilepsy, seizure frequency, and anti-seizure drug treatments. In addition, it did not correlate between the two structures and was reduced when the inflammatory status was averaged between the two structures. We also show that brain tissue not frozen within minutes after resection had significantly decreased housekeeping gene transcript levels, precluding the possibility of using post-mortem tissues to assess physiological baseline transcript levels in the hippocampus. We thus used rat models of mTLE, induced by status epilepticus (SE), that have the advantage of providing access to physiological baseline values. They indisputably indicated that inflammation measured during the chronic phase of epilepsy was much lower than the explosive inflammation occurring after SE, and was only detected when epilepsy was associated with massive neurodegeneration and gliosis. Comparison between the inter-individual variability measured in patients and that established in all epileptic and control rats suggests that some mTLE patients may have very low inflammation in the hippocampus, close to control values. However, the observation of elevated inflammation in the amygdala of some patients indicates that inflammation should be studied not only at the epileptic hippocampus, but also in the associated brain structures in order to have a more integrated view of the degree of inflammation present in brain networks involved in mesial temporal lobe epilepsy.

## 1 Introduction

Considerable research attention has been directed towards a role for neuroinflammation as one of the primary drivers of epileptogenesis occurring after brain insults and as a self-perpetuating factor of epileptic seizure activity [24, 35, 42, 46, 48, 49]. Elevated concentrations of inflammatory markers, e.g. pro-inflammatory cytokines IL1β, IL6 and TNF and chemokines, have been measured in cerebrospinal fluid and serum of patients that suffered various epileptogenic brain insults [22], but also in different forms of epilepsy [27, 52]. Access to surgically resected tissue in mTLE patients allowed evaluation of inflammatory status within the epileptic focus. Studies in human brain tissue evaluated the expression levels of certain inflammation markers in resected hippocampus of mTLE patients [22, 23, 52]. They all revealed a particularly high pro-inflammatory state in the hippocampus of mTLE patients. However, all these studies, even if they present comparisons with non-epileptic tissue, suffer from the absence of control tissues collected under conditions similar to those of operated mTLE patients. Control tissues are often autopsy specimen from people with no history of epilepsy or brain-related disease and who died without associated brain damage. Furthermore, when mentioned, sampling times range from 4 to 20.5 hours post-mortem, which is significantly longer than surgical collection of tissue from mTLE patients, with samples usually managed immediately, either by freezing [2, 10, 21, 31, 41] or by fixation [2, 10, 15, 23, 36, 40].

In the large number of studies that have been carried out over the past 2 decades, the gene markers of inflammation were measured at the level of mRNAs or proteins, by methods today recognized as very little, if at all, quantitative. Only two studies have recently reported the quantitative evaluation of some inflammatory markers from early epileptogenesis to epilepsy onset in rodent models of epilepsy [7, 18].

In our study, after demonstrating that mRNAs of three housekeeping genes were rapidly degraded in the minutes / hours following the surgical resection of the hippocampus when the resected tissues were not immediately frozen in liquid nitrogen, the first objective was to evaluate, using calibrated reverse transcription and quantitative PCR, the dispersion of the mRNA levels of prototypical inflammatory markers measured in 22 patients with drug-resistant mesial temporal lobe epilepsy who underwent epilepsy surgery, including resection of the hippocampus and the amygdala. Then, in the absence of appropriate human control tissue, we modeled mTLE in rats, which allowed us to assess not only the physiological baseline levels of inflammatory markers in the hippocampus, but also the time course of the inflammatory response during epileptogenesis and in the long term after the onset of epilepsy. Quantitative RNAscope *in situ* hybridization studies have been performed to identify cells that express IL1β gene throughout this time course.

## 2 Material and Methods

### Study Design

#### Study 1

Impact of delayed cryopreservation in the processing of human brain samples on mRNA levels of housekeeping genes (HSKG) determined by RT-qPCR. Three consecutive groups have been constituted. In groups #1 (n=13) and #3 (n=9), samples used for RT-qPCR were immediately frozen in liquid nitrogen after resection. In group #2 (n=10), freezing of the samples was 45-90 min delayed, as detailed below.

#### Study 2

Evaluating transcript levels of inflammatory markers in the resected hippocampus and amygdala of mTLE patients using RT-qPCR. After quantification of 3 genes unrelated to the inflammatory cascade in Study 1, the samples of patients included in the group #1 and 3 mentioned just above have been selected (n=22) for this experiment.

#### Study 3

Contribution of blood cells contained within capillaries of non-perfused brains to the levels of inflammatory markers measured in the hippocampus. Status epilepticus (SE) was induced at postnatal day (P) 42 (P42) by pilocarpine (Pilo-SE), and both rats subjected to SE and control rats were killed 7 hours after SE. At termination time, brains were collected from rats that were transcardially perfused with saline (control rats: n=5; SE rats: n=4) or not (control rats: n=5; SE rats: n=5).

#### Study 4

Evaluation of gene expression at transcript level in the rat hippocampus during epileptogenesis and chronic epilepsy. Pilo-SE was induced in weanlings (W) at P21 or juvenile (J) rats at P42. Hippocampus of rats were dissected after transcardial perfusion of NaCl and the inflammatory profile was evaluated by RT-qPCR. Analysis was performed in rats sacrificed at different time points after SE: during epileptogenesis, that is at 7 hours (W, n=7; J, n=6), 1 day (W, n=8; J, n= 6), 9 days (W, n=10; J, n=7) post-SE, and once chronic epilepsy was developed in all rats, i.e. 7 weeks post-SE (W, n=8; J, n=8). Brains of control rats were also collected; however, to reduce the number of animals used, some time points have been pooled: W rats (7h, 1 day and 9 days: n=10; 7 weeks: n=6) and J rats (7h and 1-9 days: n=6; 7 weeks: n=6).

#### Study 5

Astroglial and microglial activations evaluated using GFAP- and ITGAM-immunofluorescent detections, respectively, in the rat hippocampus at 1 day (W, n=4; J, n=5), 9 days (W, n=6; J, n=7) and 7 weeks post-SE (W, n=6; J, n=7), induced in W and J rats, and in respective controls (W, n=3 for 1-9 days, n=5 for 7 weeks; J, n=5 for both 1-9 days and 7 weeks).

#### Study 6

Distribution and quantitation of IL-1β transcript were evaluated using RNAscope®-based quantitative *in situ* hybridization in the hippocampus of 5 patients with mTLE (3 with high and 2 with low tissue levels of IL-1β mRNA determined by RT-qPCR) and of rats subjected to Pilo-SE at P42 and sacrificed 7 hours (n=3), 1 day (n=5), 9 days (n=3) and 7 weeks (n=3) post-SE and in respective controls (7h and 1-9 days: n=2; 7 weeks: n=2).

### Patients

Human brain tissues were obtained from 32 patients with drug-resistant mesial temporal lobe epilepsy (mTLE) who underwent anterior temporal lobectomy at the Epilepsy Department of the Lyon’s University Hospital, France, between 2009 and 2012. Evaluation of eligibility for epilepsy surgery, including presurgical work-up, was performed as described elsewhere [38]. Pre-operative written informed consent was obtained from all patients for the use of resected brain tissue for research purpose.

The first group of patients included 6 men (15-56 years) and 7 women (15-51 years); the second included 7 men (19-50 years) and 3 women (17-37 years); the third included 4 men (14-49 years) and 5 women (12-42 years). The detailed clinical data of each patient are listed in Table 1.

**Table 1.**
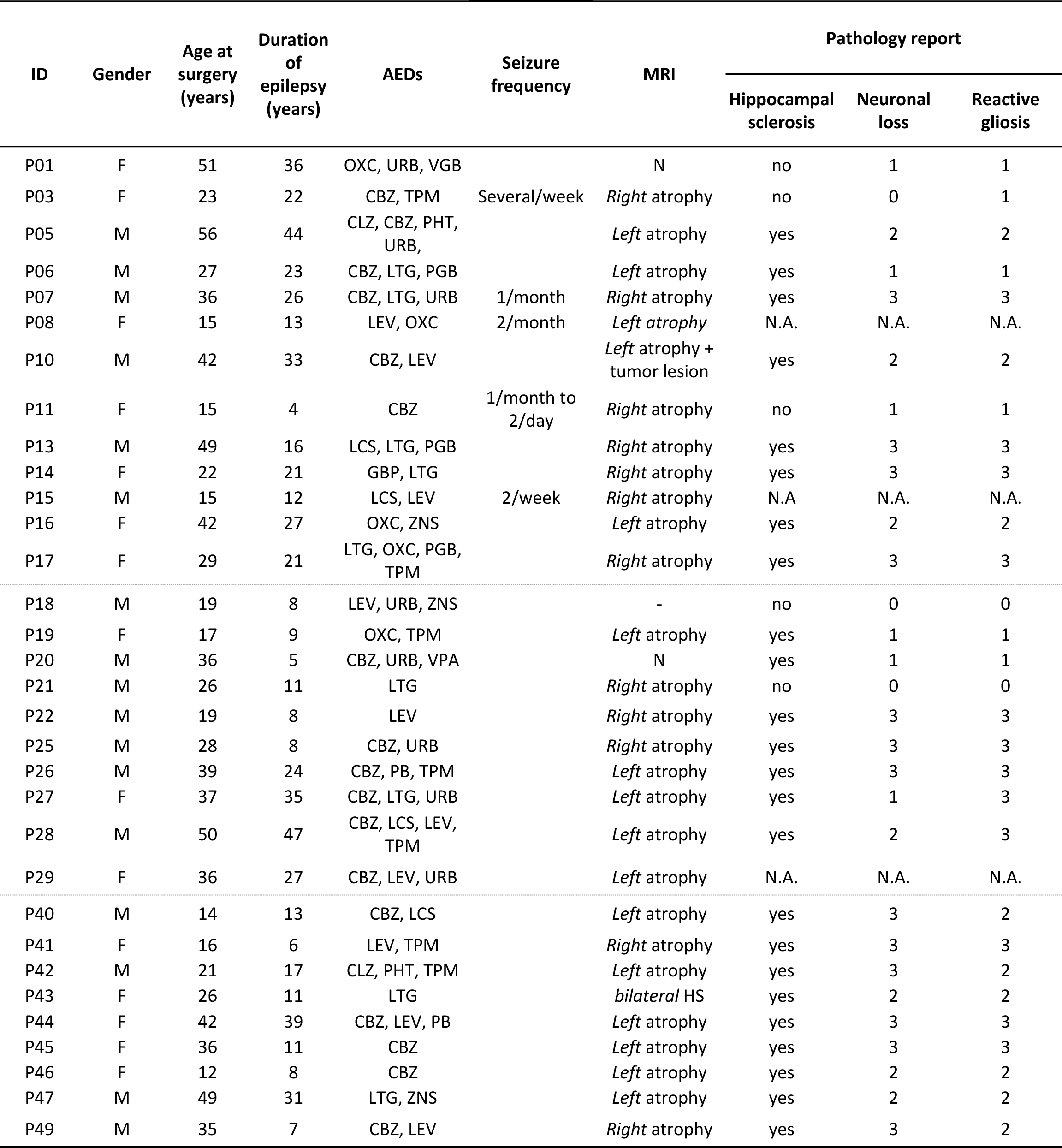
Clinical characteristics of TLE patients from group 1 (G1: P01-P17), group 2 (G2: P18-P29) and group 3 (G3: P40-P49). Data not shown in the table were not available in the patients’ medical record. Neuronal loss and reactive gliosis scoring: 0: not present; 1: mild; 2: moderate; 3: severe. Abbreviations: N: normal; HS: *hippocampal sclerosis*; AB: amyloïd bodies; N.A.: pathology report not available; NL: neuronal loss; O: oedema; RG: reactive gliosis. Anti-epileptic drugs (AEDs): CLZ: clonazepam; CBZ: carbamazepine; GBP: gabapentin; LEV: levetiracetam; LCS: lacosamide; LTG: lamotrigine; OXC: oxcarbazepine; PB: phenobarbital; PHT: phenytoin; PGB: pregabalin; TPM: topiramate; URB: urbanil; VGB: vigabatrin; VPA: valproate; ZNS: zonisamide

### Collection of surgical specimen

Hippocampi and amygdala were resected *en bloc* by the neurosurgeon (MG) and immediately given to an investigator in charge of prepation of the specimen in the operating romm. They were rinsed for 1 min in ice-cold saline and cut in 3 equal parts (for the hippocampi, cuts were performed perpendicularly to the longitudinal axis from the head to the body): the first part was immediately frozen in liquid nitrogen for 22 samples and then stored at −80°C or immersed into an ice-cold RNAlater® solution for 45 to 90 min before freezing in liquid nitrogen; the second part was fixed for 72 hours in an ice-cold 4% paraformaldehyde solution, immediately after resection (n=22) or after a 45-90 min delay (n=10), cryoprotected into an ice-cold 30% sucrose solution prepared in 0.1M phosphate buffer, frozen at −40°C in isopentane and then stored at −80°C; and the third part was used for routine histopathological evaluation.

### Animals

All animal procedures were in compliance with the guidelines of the European Union (directive 2010-63), taken in the French law (decree 2013/118) regulating animal experimentation, and have been approved by the ethical committee of the Claude Bernard Lyon 1 University (protocol # BH-2008-11). We used a tissue collection bank generated by TIGER team in 2009-2012. Briefly, male Sprague-Dawley rats (Harlan/Envigo, The Netherlands) were used in these experiments. They were housed in a temperature-controlled room (23 ± 1°C) under diurnal lighting conditions (lights on from 6 a.m to 6 p.m). Pups arrived at 15 day-old and were maintained in groups of 10 with their foster mother until P21. Beyond that age, rats were maintained in groups of 5 in 1,800 cm^2^ plastic cages, with free access to food and water. After SE, rats were weighed daily until they gained weight.

### Pilocarpine-induced *status epilepticus* (SE)

SE was induced by pilocarpine, injected at day 21 or 42. To prevent peripheral cholinergic side effects, scopolamine methylnitrate (1 mg/kg in saline, s.c.; Sigma-Aldrich) was administered 30 min before pilocarpine hydrochloride (25 mg/kg at P21 and 350 mg/kg at P42, in saline, i.p.; Sigma-Aldrich). For P21 rat pups, lithium chloride (127 mg/kg in saline, i.p.; Sigma-Aldrich) was injected 18 hours before scopolamine. After 30 min of continuous behavioral SE at P21 and 2 hours at P42, 10 mg/kg diazepam (i.p.; Valium; Roche®) was injected, followed, 90 min later for P21 and 60 min later for P42, by a second injection of 5 mg/kg diazepam to terminate behavioral seizures. Control rats received systematically equivalent volumes of saline solution. The animals were then sacrificed at various time points: 7 hours, 1 day, 9 days and 7 weeks after SE.

### Animal care after Pilo-SE

Control and treated rats were weighted every day during the first two weeks following Pilo-SE, and then every week until termination of the experiment. Daily abdominal massages were performed twice a day during the first week to activate intestinal motility, which was disrupted following Pilo-SE.

### Detection of spontaneous recurrent seizures (SRS)

Electroencephalographic recordings were excluded to determine epilepsy onset due to pilot experiments that showed that the sole implantation of screws into the skull induced significant and lasting inflammation over time in the cortex and, to a lesser extent, in the hippocampus. As a result, epilepsy onset was determined according to clinical criteria.

As previously reported [12, 25], development of a chronic epileptic state, i.e. with SRS, was confirmed in all rats subjected to Pilo-SE at P42 by the end of the 2^nd^ week post-SE. However, there is an inconsistency in the literature about the proportion of rats that develop SRS and the time of SRS onset when Pilo-SE is induced at P21 in Sprague-Dawley rats [12, 32]. We thus induced Pilo-SE in a group of 7 male rats at P21 and determined whether they developed SRS by the 7^th^ week post-SE, using a video-EEG monitoring performed 2-3 days a week, for 24 consecutive hours each time, from the 4^th^ to the 7^th^ week post-SE. When rats were placed in their recording cage, they were each time subjected to a handling-induced seizure (HIS) test, which consisted in restraining rats for 10 seconds at the level of the chest with gentle pressure. During the 3^rd^ week post-SE, rats were implanted under 3% isoflurane anesthesia with three screw electrodes positioned over the frontal and parietal cortices, and over the cerebellum used as ground electrode. Electrodes were connected to a multipin socket and secured to the skull using a thin layer of dental adhesive (Super-Bond C&B) and acrylic dental cement. One week after surgery, rats started to be connected to the video-EEG setup. All rats had developed HIS by the end of the 5^th^ week post-SE and all had SRS at the end of the 6^th^ week post-SE. During the 7^th^ week post-SE, the average number of seizures was 1.1 ± 0.3 seizure/day (n=7). Based on this result, for rats subjected to Pilo-SE and used for inflammation analysis, epilepsy development was monitored by testing the occurrence of HIS three times a day from the 5^th^ week post-SE. Once HIS were developed on 2 consecutive trials, rats were observed by experimenters for 5 consecutive hours (between 01:00 and 06:00 p.m) over the following days to detect the presence or absence of SRS. All rats were declared as “epileptic” (EPI) by the end of the 6^th^ week post-SE.

### Ex Vivo Procedures

All rats were deeply anesthetized with a lethal dose of pentobarbital (100 mg/kg; Dolethal) before being sacrificed. All rats were transcardially perfused with sodium chloride (NaCl 0.9%) for 3 min at 30 mL/min. For RT-qPCR analysis, hippocampus were rapidly microdissected, frozen in liquid nitrogen, and stored at −80°C. For immunochemistry analysis, animals were transcardially perfused (30 mL/min) with 4% paraformaldehyde in 0.1 M phosphate buffer. After cryoprotection in 30% sucrose, brains were frozen at −40°C in isopentane and stored at −80°C.

### RNA extraction and quantification of transcript level variations by reverse transcriptase real-time polymerase chain reaction (RT-qPCR)

Brain structures frozen in liquid nitrogen were crushed using Tissue-Lyser (Qiagen®) in 250 µL of ultrapure RNase-free water (Eurobio). Nucleic acids were extracted by adding 750 μL Tri-Reagent LS (TS120, Euromedex) and 200 μL chloroform (VWR®). After precipitation with isopropanol (I-9516, Sigma-Aldrich®), washing in 75% ethanol (VWR) and drying, total nucleic acids were resuspended in 50 μL ultrapure water and treated with DNAse I (Turbo DNA Free® kit; AM1907, Ambion®) to eliminate any trace of possible genomic DNA contamination. The purified total RNAs were then washed using the RNeasy® minikit (Qiagen®) kit. After elution, the total RNA concentration was determined for each sample on BioDrop® µLite. The quality of total RNAs was verified on microgel chips using LabChip® 90 (Caliper), which provides an RNA Integrity Number (RIN) value by analyzing the integrity of two ribosomal RNAs (18S and 28S) predominantly present in all tissue RNA extracts. All selected samples had a RIN value greater than 7.0, and were stored at −80°C until use. Total tissue RNAs (480 ng) were reverse transcribed to complementary DNA (cDNA) using both oligo dT and random primers with PrimeScript RT Reagent Kit (Takara) according to manufacturer’s instructions, in a total volume of 10 µL. In RT reaction, 300 000 copies of a synthetic external non-homologous poly(A) standard messenger RNA (SmRNA; [1], patent WO2004.092414) were added to normalize the RT step [39]. cDNA was diluted 1:13 with nuclease free Eurobio water and stored at −20°C until further use. Each cDNA of interest was amplified using 5 µL of the diluted RT reaction by the “real-time” quantitative polymerase chain reaction (PCR) technique, using the Rotor-Gene Q thermocycler (Qiagen®), the SYBR Green Rotor-Gene PCR kit (Qiagen®) and oligonucleotide primers specific to the targeted cDNA. The sequences of the specific forward and reverse primer pairs were constructed using the Primer-BLAST tool or using the “Universal Probe Library” software (Roche Diagnostics). Sequences of the different primer pairs used are listed in Supplementary Table S1 for humans and Supplementary Table S2 for rats. The number of copies of each targeted cDNA contained in 5 µL of the diluted RT reaction was quantified using a calibration curve based on cascade dilutions of a solution containing a known number of cDNA copies.

**Table 2.** Number of cDNA copies (mean ± SEM) after reverse transcription in the resected hippocampus of patients with refractory epilepsy (n = 22). P41 and P49 patients have been chosen for exemplification, as they present with the lowest and the highest values of the pro-inflammatory index (Figure 3), respectively

| cDNA copies number in hippocampus of epileptic patients |  |  |  |  |  |
| --- | --- | --- | --- | --- | --- |
| | Mean ± SEM | Min | Max | P41 | P49 |
| **IL1β** | 28 245 ± 5 388 | 3 474 | 114 957 | 7 833 | 56 674 |
| **IL6** | 948 ± 327 | 0 | 5 634 | 75 | 5 634 |
| **TNF** | 3 727 ± 1 115 | 0 | 20 485 | 0 | 20 485 |
| **MCP1** | 201 347 ± 26 979 | 37 289 | 425 797 | 37 289 | 415 250 |
| **MIP1α** | 99 659 ± 16 512 | 13 683 | 278 864 | 13 683 | 178 990 |
| **IL10** | 425 ± 82 | 0 | 1274 | 0 | 1 274 |
| **GFAP** | 4 274 122 ± 481 786 | 736 298 | 7 849 435 | 1 190 100 | 7 849 435 |
| **ITGAM** | 1 661 ± 279 | 99 | 4 314 | 181 | 4 024 |

Pro-inflammatory (PI-I), anti-inflammatory (AI-I), inflammation cell (IC-I) and housekeeping gene (HSKG-I) indexes were calculated for each series of individuals to be compared using a specific set of genes: IL1β, IL6, TNF, MCP1 and MIP1α for PI-I; IL4, IL10 and IL13 for AI-I; ITGAM and GFAP for IC-I; DMD, HPRT1 and GAPDH for HSKG-I. For each individual, the number of copies of each transcript has been expressed in percent of the averaged number of copies measured in the whole considered population of individuals. Once each transcript is expressed in percent, an index is calculated by adding the percent of each transcript involved in the composition of the index and expressed in arbitrary units (A.U.). Each time that an index is presented, the groups of individuals constituting the population is specified.

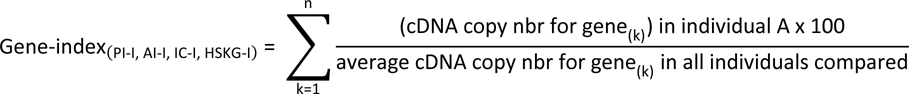

### Tissue processing for histological procedures

Cryostat-cut (40 µm thick) sections from mTLE patient tissue samples or from rat samples were transferred into a cryopreservative solution composed of 19.5 mM NaH_2_PO_4_.2H_2_O, 19.2 mM NaOH, 30% (v/v) glycerol and 30% (v/v) ethyleneglycol and stored at −25°C.

### Immunohistochemistry

For immunohistofluorescent detections, free-floating sections (40 μm thick) from paraformaldehyde-fixed tissue were incubated with a rabbit polyclonal anti-GFAP antibody (1:1,000; AB5804, Chemicon) to label astrocytes, a mouse monoclonal anti-ITGAM antibody (1:1,000; CBL1512Z, Chemicon) to detect microglia and immunocompetent cells, a goat polyclonal anti-CD14 antibody (1:1,000; sc-5749, Santa-Cruz) to detect monocytes, a mouse monoclonal anti-NeuN antibody (1:1,000; MAB-377, Chemicon) to label neurons, and finally rabbit polyclonal anti-IL1β antibody (1:200; #250716, Abbiotec). Some sections were also incubated with a combination of a mouse monoclonal anti-GFAP antibody (1:1,000; G3893, Sigma-Aldrich) and the above-described polyclonal anti-GFAP antibody to verify whether both antibodies provided similar detections of GFAP protein. Fluorescent secondary antibodies (Alexa-Fluor-conjugated antibodies; Molecular Probes) used were: A488-donkey anti-rabbit IgG antibody (1:1,000; A-21206), A647-donkey anti-mouse IgG antibody (1:1,000; A-31571), A633-goat anti-mouse IgG antibody (1:500; A-21052), A633-donkey anti-goat IgG antibody (1:1,000; A-21082). Sections were then mounted on SuperFrost Plus slides and coverglassed with Prolong Diamond Antifade reagent (Molecular Probes). Dual-immunolabelings of GFAP and ITGAM were observed using a Carl Zeiss Axio Scan.Z1 Digital Slide Scanner with a resolution of x40. Dual-immunolabelings of GFAP, and single immunolabeling of either NeuN or IL1β were observed using a LSM800 confocal microscopy system (Zeiss) with ZEN imaging software (Zeiss). Images were then imported into Adobe Photoshop CS6 13.0 (Adobe Systems) for further editing. All sections were analyzed under identical conditions of photomultiplier gain, offset and pinhole aperture, allowing the comparison of fluorescence intensity between regions of interest. For NeuN, ImageJ software was used to measure surface areas of fluorescence using thresholding procedures. The quantification of the immunofluorescent surface area was performed on stacks of 12 images taken over a thickness of 11.36 µm with a step of 1.03 µm. For colorimetric immunohistodetection of NeuN, free-floating sections were sequentially incubated with the mouse monoclonal anti-NeuN antibody (1:1,000; MAB-377, Chemicon), and with a biotinylated donkey anti-mouse IgG (1:1,000; Jackson Immuno Research, 715-065-151), and revealed by the Avid-Biotin Complex (ABC)-peroxydase (1:1,000; Vector, PK-6100) in presence of DAB.

### *In Situ* Hybridization using RNAscope®

Probes were designed by ACD (Advanced Cell Diagnostics, Newark, New Jersey) to hybridize to IL1β, ITGAM and GFAP mRNA molecules with species specificity (*Homo sapiens* -Hs-probes for humans; *Rattus Norvegicus* -Rn-for rats). The RNAscope® Multiplex Fluorescent Reagent Kit v2 (Cat. 323100) and the hybridization oven (HybEZ Oven) were also obtained from ACD. The RNAscope® assay was performed as described by the supplier. Briefly, the staining protocol included five steps: pretreatment with protease, hybridization of target probes, amplification of the signal, detection of the signal and mounting of the slides.

Selected tissue sections of resected hippocampus from mTLE patients or selected rat tissue section including the hippocampus were removed from cryoprotectant solution and rinsed in phosphate-buffered saline (PBS) three times. RNAscope® assays were performed on tissue mounted on SuperFrost slides. Sections went through treatment with Protease III solution during 30 minutes at 40°C. Three different probes were then used to localize mRNAs of IL1β (Hs-IL1β, Cat. 310361; Rn-IL1β, Cat. 314011), ITGAM (Hs-ITGAM, Cat. 555091-C3; Rn-ITGAM, Cat. 300031-C3) and GFAP (Hs-GFAP, Cat. 311801-C2; Rn-GFAP, Cat. 407881-C2). Sections subsequently passed through amplification steps followed by fluorescent labeling in Opal 520, Opal 570 and Opal 690 (NEL810001KT, PerkinElmer) at 1:1000 dilution with amplification diluent. Sections were then counterstained with DAPI and coverglassed with Prolong Diamond Antifade reagent (Molecular Probes). Slides were observed using a TCS SP5X confocal microscopy system (Leica). All sections were analyzed under identical conditions of photomultiplier gain, offset and pinhole aperture, allowing the comparison of fluorescence intensity between regions of interest. Then, for each of the hybridized probe, ImageJ software was used to measure areas of fluorescence using thresholding procedure.

### Data and statistical analysis

GraphPad Prism (v.7) software was used to statistically analyze data. Majority of data are expressed as mean ± SEM of the different variables analyzed. Transcript levels are also expressed using box-and-whisker plots to illustrate the distribution of the considered cohort. Statistical significance for within-group comparisons was calculated by one-way or two-way analysis of variance (ANOVA) with Bonferroni or Tukey’s *post hoc* test. The p value of 0.05 defined the significance cut-off. Correlations were assessed using Spearman’s rank correlation test.

## 3 Results

### Human brain tissues with delayed cryopreservation are not suitable controls for transcriptomic studies

The main clinical characteristics of the 32 patients included in the study are summarized in Table 1. Ideally, surgically resected brain tissues should be frozen at a very low temperature immediately after collection, so as to preserve the molecules to be measured. These conditions were those observed for Patient Groups 1 and 3 (G1 and G3), whose resected brain tissues were frozen less than 5 minutes after neurosurgical removal. For logistic reasons, it was temporarily decided to delay the freezing procedure of the resected tissues from Patient Group 2 (G2). To this end, tissues were transferred into ice-cold RNALater® immediately after their resection, then given to the research staff in charge of freezing them in liquid nitrogen back to the laboratory within a time interval ranging between 45 and 90 minutes. RNALater® has been developed to preserve RNA integrity even if samples are stored for days to weeks at 4°C after collection either before freezing or direct extraction of total RNAs [17]. We first compared the mRNA levels of three housekeeping genes (HSKG = GAPDH, HPRT1 and DMD) between G1 (P01-P17, n=13) and G2 (P18-P29, n=10). To ensure mRNA quantification independent of any internal control (i.e. by a so-called invariant gene), we used an external standard mRNA (the SmRNA patented by our group) [1].

A very large decrease was observed in G2 (7.2-fold less than G1 for DMD: p=0.0022; 4.7-fold less than G1 for GAPDH: p=0.0371; 14-fold less than G1 for HPRT1: p=0.0008, Fig. S1). By switching back to the first freezing protocol (i.e. freezing immediately after resection) for G3 (P40-P49, n=9), the average values for the 3 housekeeping genes were closer to that obtained for G1 (Fig. 1). Significant differences were observed between G1 and G3 for GAPDH and HPRT1 transcripts. Although one can only speculate at this point on the interpretation of this result, one explanation could be that the transcript levels of these genes are so variable from one patient to another that it may be that when these patients are randomly assigned between two groups, these differences become significant. All of the samples used in this study had RIN values >7, attesting that all RNA samples were of excellent quality, according to the integrity of 18S and 28S ribosomal RNAs. Overall, these results indicate that delayed cryopreservation protocol caused alteration of the transcript levels in human resected hippocampi, precluding the use of autopsy/post-mortem tissue as valid controls, in particular to determine reference / basal levels of neuroinflammatory markers. For the rest of the study, values of patient group 2 were excluded.

**Fig. 1.**
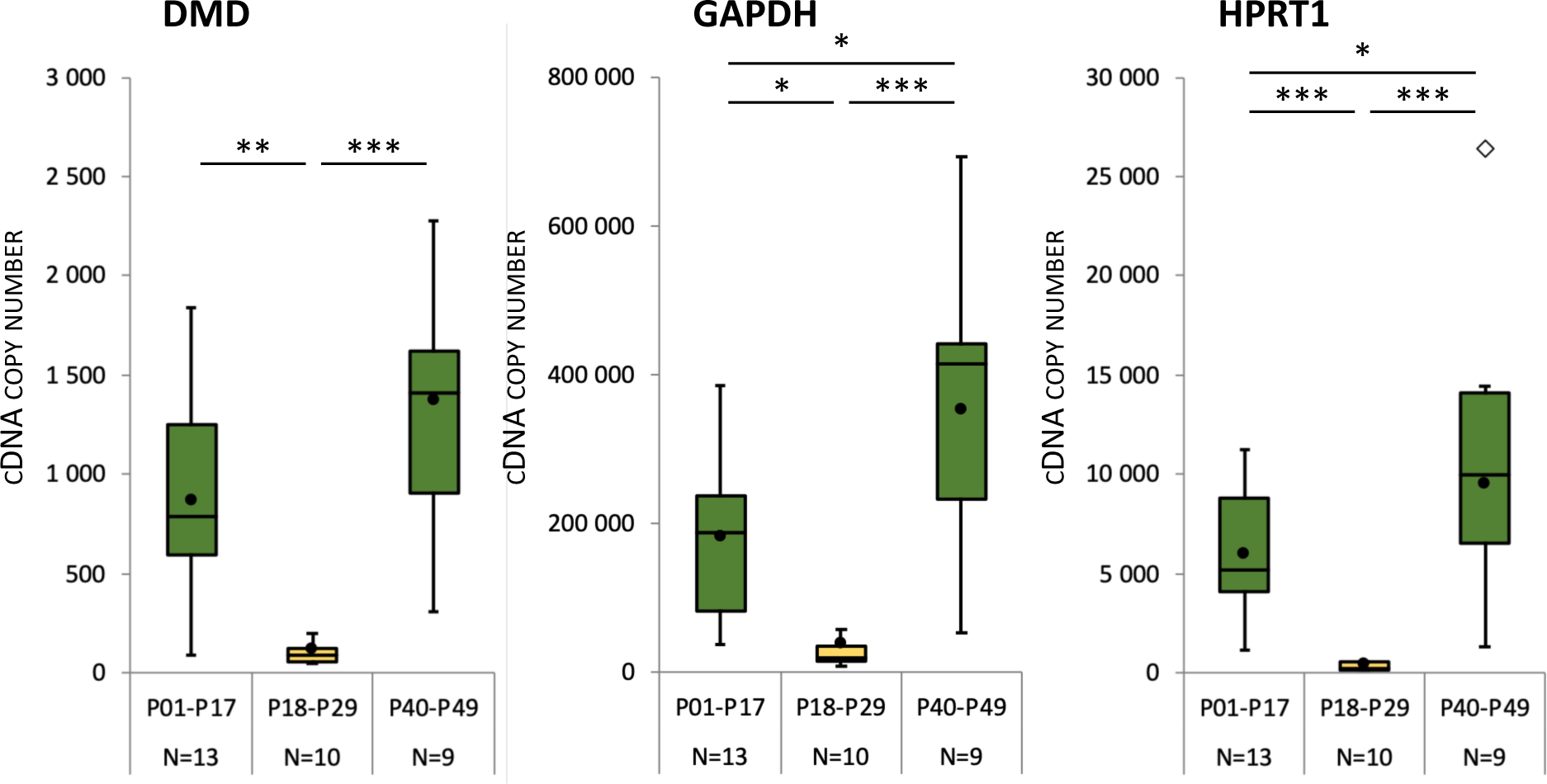
Delayed cryopreservation of human brain tissue significantly alters transcript levels of housekeeping genes. Hippocampus from 3 groups of TLE patients were resected surgically and frozen in liquid nitrogen during the 5 minutes (Group 1: P01-P17; Group 3: P40-P49; green box and whisker plots) or 45 to 90 minutes (Group 2; P18-P29 group, yellow box and whiskers plot) after resection. Transcript levels of three housekeeping genes (DMD, GAPDH and HPRT1) were quantified. Box-and-whisker plots model the distribution of each value around the median of the cDNA copy number measured by RT-qPCR. Mean is represented by black dots. Outliers are represented by diamonds. Tukey’s *post-hoc* analysis following one-way ANOVA: * *p*<0.05, ** *p*<0.01, *** *p*<0.001

### Inflammation in the hippocampus of mTLE patients is highly variable

Twenty-two fresh frozen surgically resected hippocampi of mTLE patients (P01-P17 and P40-49 from G1 and G3; 12F, 10M, 31 ± 14 years) were subjected to gene-specific transcript quantification for a set of 11 inflammatory markers including pro-inflammatory (IL1β, IL6, TNF, IFNγ) and anti-inflammatory (IL4, IL10, IL13) cytokines, chemokines (MCP1, MIP1α) and cell markers (microglia/macrophages: ITGAM, astrocytes: GFAP). In our relatively small-scale gene expression analysis study, we measured transcripts by calibrated RT and qPCR rather than the corresponding proteins by immunohistofluorescence, western blot or Elisa, for example. Indeed, the latter methods depend on the ability of the antibodies used to recognize the targeted proteins, which may not be equivalent and thus generate interpretation biases, as shown by the detection of the GFAP protein on brain sections using two antibodies directed against the same protein (Fig. S1). For each patient, all the above-mentioned gene transcripts could be detected and then quantified, except those of IFNγ, IL4 and IL13 that were not detected in any of the 22 samples, even when using several primer pairs designed in different parts of the corresponding cDNAs. Individual cDNA value for each marker was expressed in percent of the calculated average (n=22) value (Fig. 2). IL6 and MCP1 had the highest and lowest interindividual variability, respectively.

**Fig. 2.**
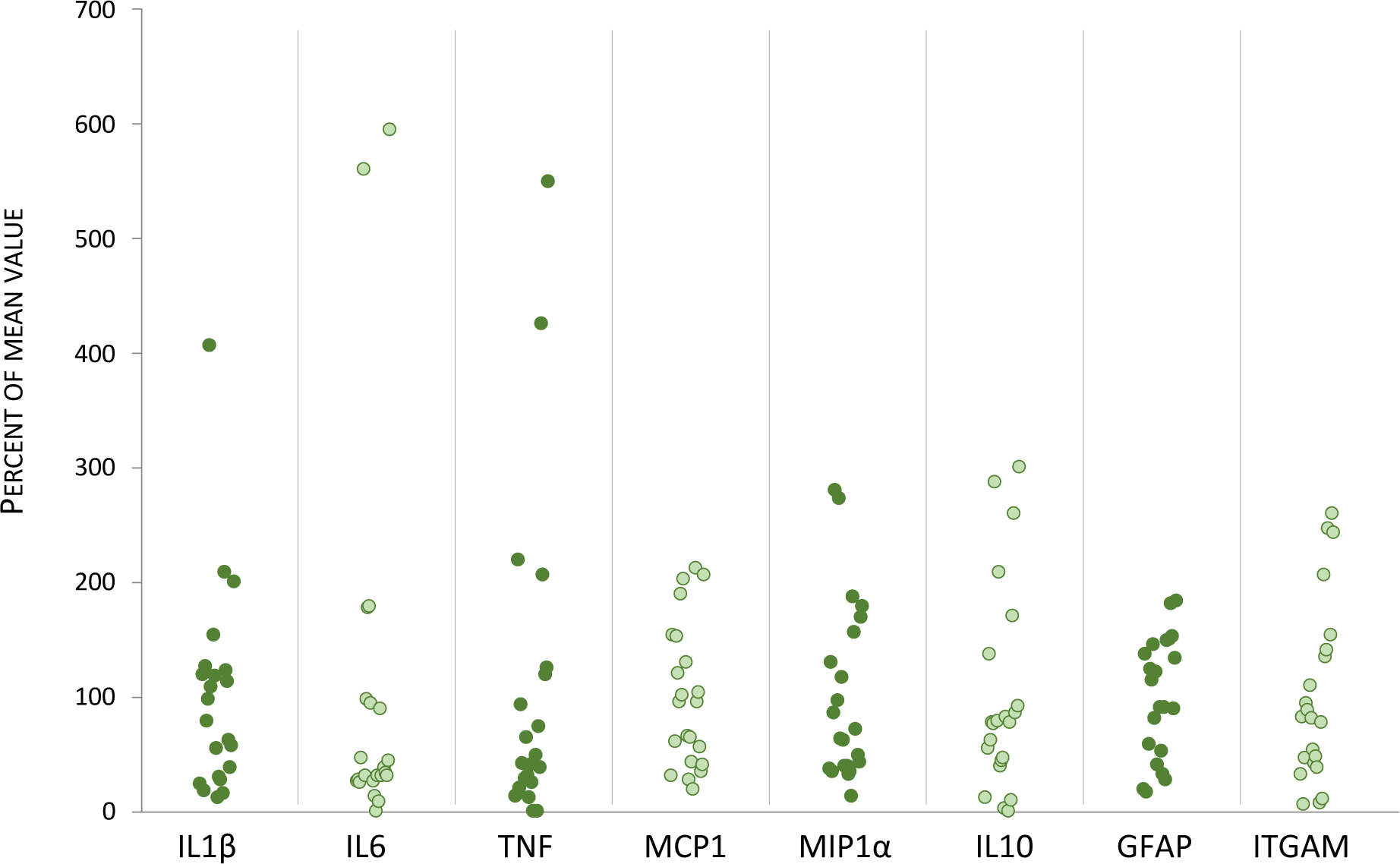
Patients with TLE are heterogeneously distributed regarding the molecular and cellular markers of inflammation measured in the hippocampus. Transcript level of pro-inflammatory cytokines (IL1β, IL6, TNF), chemokines (MCP1, MIP1⍺), anti-inflammatory cytokine IL10, and cellular markers (GFAP for astrocytes, ITGAM for microglia/macrophages) were measured in resected hippocampus of TLE patients (n=22). Each point represents a patient, and individual values are expressed in percent of the mean value for each marker

Not all the lowest values are observed in the same patient, nor are the highest values (Table 2). For example, patient P41 who had the lowest values for TNF, MCP1 and MIP1α did not have the lowest values for IL1β and IL6. Similarly, patient P49, who had the highest values for TNF and IL6, did not have the highest values for IL1β, MCP1 and MIP1α (Tables 2 and S3). Therefore, to provide a general overview of the inflammatory status for each patient, we calculated a pro-inflammatory index (PI-I) and an inflammation cell index (IC-I) that integrate for each patient the average normalized expression of each individual cytokine/chemokine or each individual cell marker, respectively. Patients P41 and P49 had the lowest (56 A.U.) and the greatest (1,730 A.U.) pro-inflammatory index, respectively (Fig. 2A-B), corresponding to a ∼31-fold difference. It is to note that the pro-inflammatory index correlated with the inflammation cell index (IC-I = 0.195 x PI-I + 102.2; p<0.0023).

It is to note that the levels of the anti-inflammatory cytokine IL10 correlated with the pro-inflammatory index (PI-I = 3.015 (IL10 level) + 198; p<0.0004), suggesting that pro- and anti-inflammatory processes are subjected to a coordinate regulation in the hippocampus of mTLE patients.

To our knowledge, in order to normalize RT-PCR data, all prior studies used one or a combination of housekeeping genes considered as invariant between samples. We previously stressed the fact that high variability was also found in housekeeping genes (Fig. 1). We calculated a housekeeping gene index integrating DMD, GAPDH and HPRT1, which confirmed the high variability in the pool of the three housekeeping genes between patients, e.g. a 22-fold difference between patients 5 and 42 (Fig. S2A). We show that the housekeeping gene variability did not fit with that of the pro-inflammatory index (Fig. 3A and Fig. S2A). Hence, if housekeeping genes had been used to normalize the RT reaction, this would have led to biased results (Fig. S3A).

**Fig. 3.**
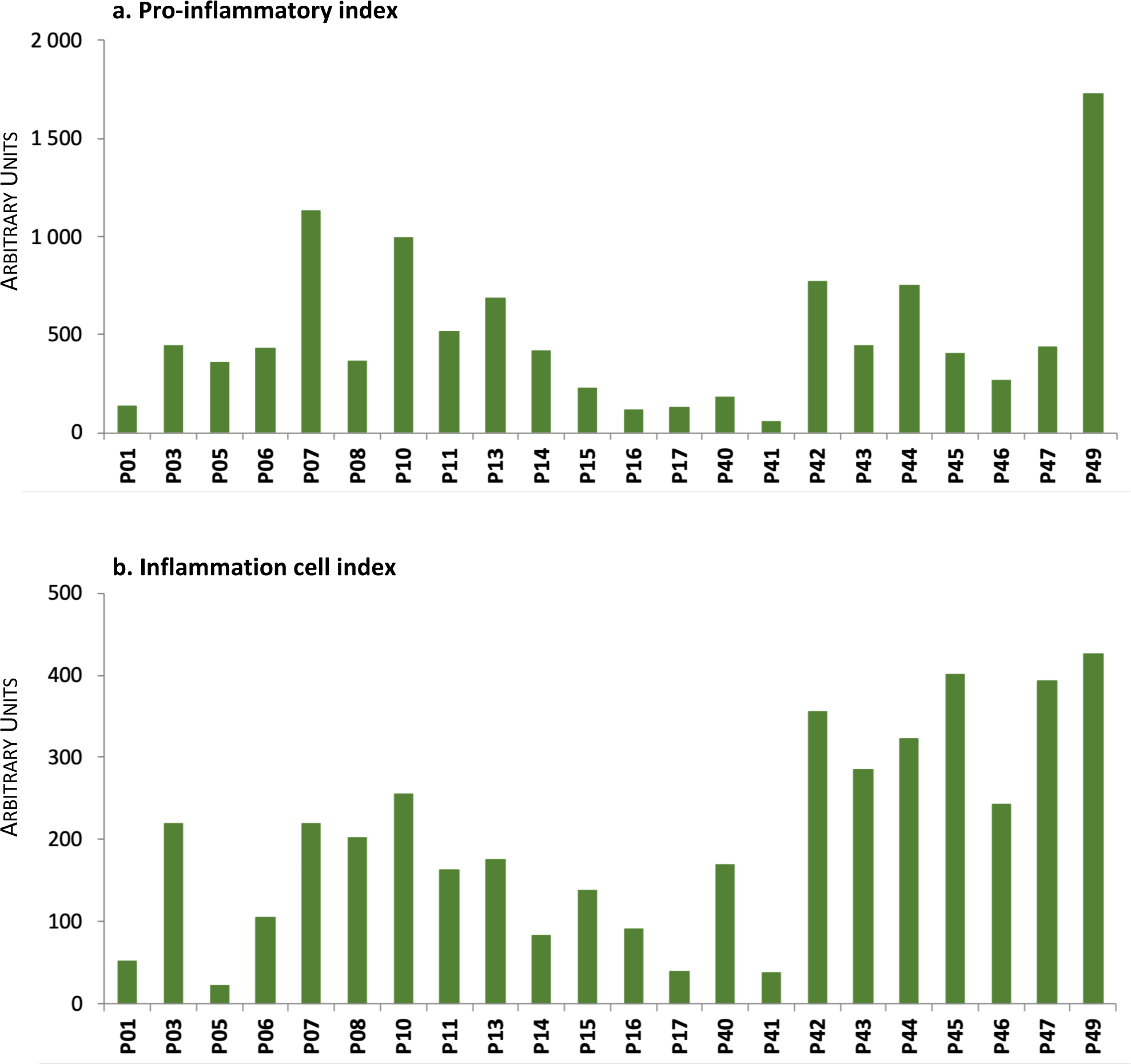
Individual inflammatory indexes in the hippocampus of TLE patients. Distribution of the values of pro-inflammatory index **(a)**, and inflammation cell index **(b)** in resected hippocampus of TLE patients (n=22). Indexes were calculated from transcript levels as described in the methods section

We next investigated whether the variation of the pro-inflammatory index was associated with relevant clinical features. The six patients with the greatest pro-inflammatory index (P07, P10, P13, P42, P44, P49) all had neuronal loss (Table 1), but neuronal loss was not systematically associated with a high pro-inflammatory and inflammation cell index values (PI-I = 106.8 x [neuronal loss score] + 27; p=0.31 and IC-I = 29.02 x [neuronal loss score] + 137.7; p=0.39).

The pro-inflammatory index (PI-I, in A.U.) did not correlate either with the age (in year) at epilepsy onset (PI-I = 14.95 x age + 342.3; r^2^=0.11939) nor with the duration (in year) of epilepsy (PI-I = −0.017 x duration + 503.5; r^2^ = 0.00002). While reports on seizure frequency before surgery were lacking for most patients, data available for 5/22 patients provide indication that rare seizures (P07, 1 seizure per month) and frequent seizures (P15, 2 seizures per week) were associated with high (1,132 A.U.) and low (228 A.U.) pro-inflammatory index values, respectively. Finally, the extent of the pro-inflammatory and inflammation cell indexes were not associated with any given anti-epileptic drug treatment (compare Table 1 and Fig. 3A-B).

Overall, our results show that some, but not all patients with refractory mTLE, had a substantial level of inflammation within the resected hippocampus. At this stage, the absence of appropriate human control tissues, as this is the case in some studies [2, 7, 33], did not allow us to know if the inflammation observed was at low or very high level. In order to provide answers to this question, the rest of this study was conducted on preclinical models in order to have access to valid control tissues and to compare the inflammatory level during chronic epilepsy to that reported during epileptogenesis [7, 18, 46].

### Circulating inflammatory markers do not contribute significantly to the quantitation performed in whole brain extracts

Resected hippocampi from patients with mTLE contain blood tissue; it was thus essential to ascertain whether the presence of blood could be a hindrance to the evaluation of brain parenchyma inflammatory status. We used rats subjected to pilocarpine-induced *status epilepticus* (SE) to evaluate the potential contribution of blood into the measures performed in brain tissue. Transcripts levels of IL1β, IL6, TNF, MCP1, MIP1α and ITGAM were compared between rats devoid of blood tissue following transcardial perfusion of sodium chloride and rats that were not subjected to perfusion (Table 3). The study was conducted in juvenile rats 7h after SE induction (perfused rats: SE-NaCl; not perfused: SE-blood) and in their respective controls (perfused rats: CTRL-NaCl; not perfused rats: CTRL-blood). Results are expressed as the percentage of the mean transcript level value measured in CTRL-NaCl group. Except for TNF, where a significant difference is observed between the two groups of controls (p <0.01), the inflammatory expression profiles are identical with or without transcardial perfusion of NaCl, showing that the level of inflammatory molecules into brain vessels remains marginal, indicating that most inflammatory molecules measured in whole brain extracts originated more from brain parenchyma than blood.

**Table 3.** Blood cells do not contribute significantly to the inflammatory markers detected in brain. Transcript level quantitation was performed in the hippocampus of control rats or epileptic rats 7h after SE, after transcardial perfusion of 0.9% NaCl or not. Brains were dissected immediately after death (CTRL-blood: n=5; SE-blood: n=5; CTRL-NaCl: n=5; SE-NaCl: n=4). NS: statistically not significant

| | IL1β | IL6 | TNF | MCP1 | MIP1α | ITGAM |
| --- | --- | --- | --- | --- | --- | --- |
| CTRL-NaCl | 100 ± 17 | 100 ± 15 | 100 ± 13 | 100 ± 55 | 100 ± 8 | 100 ± 8 |
| CTRL-blood | 140 ± 14 | 65 ± 16 | 153 ± 8 | 70 ± 8 | 90 ± 5 | 99 ± 6 |
|  | NS | NS | *p*<0.01 | NS | NS | NS |
| SE-NaCl | 2 197<br>± 794 | 23 720<br>± 10 375 | 345<br>± 100 | 131 243<br>± 23 148 | 1 151<br>± 159 | 527<br>± 66 |
| SE-blood | 1 855<br>± 757 | 24 127<br>± 13 099 | 255<br>± 32 | 124 747<br>± 19 724 | 885<br>± 155 | 617<br>± 89 |
|  | NS | NS | NS | NS | NS | NS |

### Model-specific differences in post-SE microgliosis and astrogliosis

All but one patient with mTLE demonstrated with hippocampal sclerosis on pathological examination and among them, the extent of neuronal loss and reactive gliosis was highly variable (Table 1). Therefore, to model the heterogeneity of patients with mTLE, we used two well-known rat models presenting various extents of neuronal degeneration. The first model used consisted of juvenile P42 rats subjected to pilocarpine-induced SE (Pilo-SE), characterized by extensive neuronal degeneration in the hippocampus, the piriform cortex, the amygdala and the insular agranular cortex [29, 39, 51]. By contrast, the second model used consisted of weaned P21 rats subjected to lithium-Pilo-SE, characterized by minimal or not detectable neuronal loss at 15 days post-SE or once adults [8, 9], as illustrated (Fig. 4).

**Fig. 4.**
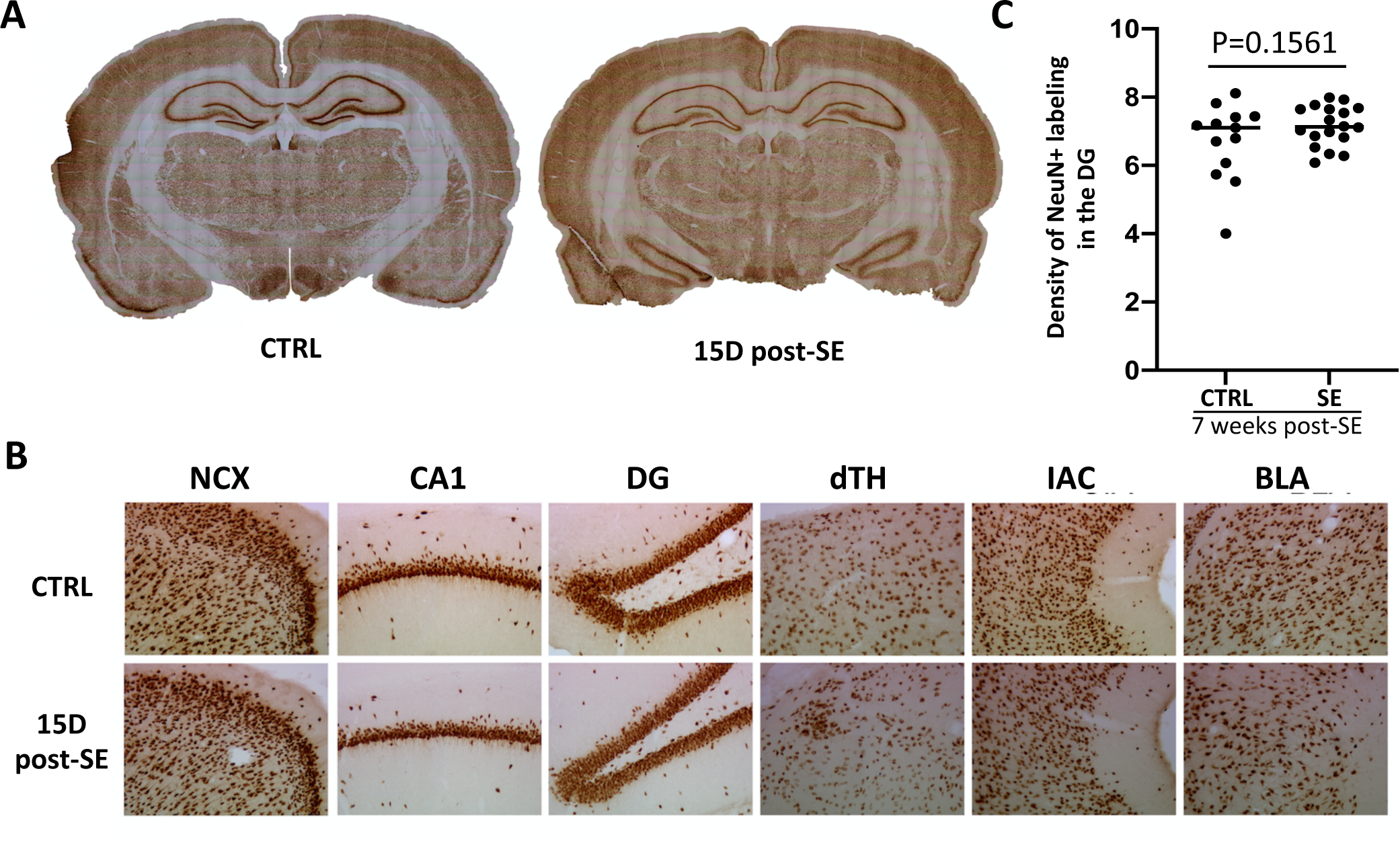
Absence of massive neurodegeneration in the brain of Sprague-Dawley rats after Pilo-SE induced at weaning. **a** Illustration of NeuN-immunolabeling at 36 days of age, in a control rat and in a rat subjected to SE 15 days earlier (P21). **b** Enlarged observations of NeuN-immunolabeling in brain regions of the sections presented in **(a)**. Note that these regions are usually affected by massive neurodegeneration when SE is induced at P42 (Nadam et al., 2007; Sanchez et al., 2009). **c** Quantification of the surface area occupied by NeuN-immunolabeling in the DG, 7 weeks post-SE (CTRL, n=13 sections from 5 rats; SE, n= 18 sections from 6 rats). Abbreviations: BLA, basolateral nucleus of the amygdala; D, day; DG, dentate gyrus; dTH, dorsal thalamus; IAC, insular agranular cortex; NCX, neocortex

In these two models, characterization of SE-induced reactive gliosis in the rat hippocampus was performed histologically during epileptogenesis (1 day and 9 days post-SE) and during the chronic phase of epilepsy (7 weeks post-SE), by double-labeling immunofluorescence targeting GFAP and ITGAM (CD11b) to evaluate astroglial and microglial/macrophage reactivity, respectively (Fig. S4). Before induction of SE, astrocytes and microglia showed low GFAP and ITGAM signal, respectively. High reactivity of both GFAP and ITGAM was observed at 1 day and 9 days post-SE in rats subjected to juvenile SE, and, to a lesser extent for rats subjected to SE at weaning. These histological results are in line with those obtained for the corresponding transcripts measured by RT-qPCR, the induction of both GFAP and ITGAM in the hippocampus of rats subjected to SE at P42 (dark blue bars) being greater during epileptogenesis to that of rats subjected to SE at P21 (light blue bars) (Fig. 5A-B). In addition, as previously reported in rats following pilocarpine-induced SE [36], preliminary experiments from our laboratory indicated that ITGAM+ round-shaped cells infiltrated the brain parenchyma beyond 7 hours and until 3 days post-SE, ther peak being observed 24 hours post-SE. These cells were identified as extravasating macrophages, as all round-shaped ITGAM+ cells in the hippocampus expressed macrophage-specific CD14 marker (Fig. 6). During the chronic phase of epilepsy, at 7 weeks after SE, GFAP and ITGAM mRNA levels decreased markedly in the hippocampus, and GFAP transcript remained higher than controls only in rats subjected to SE at P42 (Fig. 5A-B). When considering the overall markers of reactive gliosis (GFAP and ITGAM mRNAs), the inflammation cell index was always greater in rats subjected to SE at P42 compared to P21, both during epileptogenesis and the chronic phase of epilepsy (Fig. 5C).

**Fig. 5.**
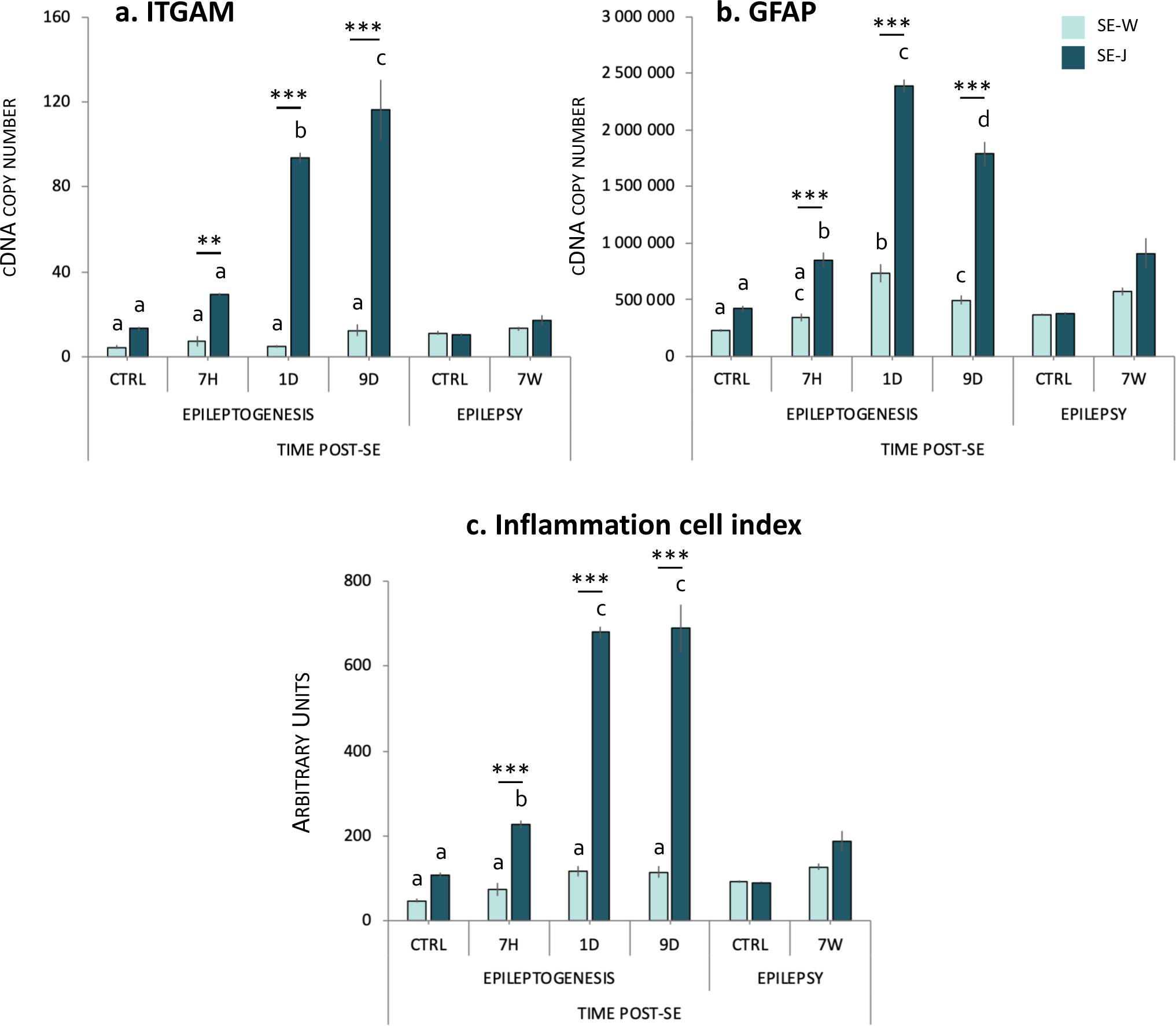
Expression of cell markers (ITGAM and GFAP) after Pilo-SE. Transcript values of ITGAM **(a)** and GFAP **(b)** and inflammation cell index **(c)** in Sprague-Dawley (SD) rats are given during epileptogenesis, i.e at 7 hours (7H), 1 day (1D), 9 days (9D) post-SE and once epilepsy was chronically installed, i.e. 7 weeks post-SE (7W) compared to respective controls. In each model (SE-W and SE-J), data from control rats have been pooled together during the epileptogenesis period (7H to 9D), after ensuring for no statistical difference between these stages. Corresponding number of copies for each gene is given in supplementary table S4. When comparing two bars within a same model, the difference is considered as statistically significant (*p*< 0.05) when letters (a, b, c, d) above the bars are different (a-b; a-c; a-d; b-c; b-d; c-d). Asterisks indicate statistical significance between the two models (SE induced at weaning or juvenile stage) at a same post-SE time. The statistical analysis only represents significant differences during epileptogenesis. 7H: SE-W, n=7; SE-J, n=6. 1D: SE-W, n=8; SE-J, n=6. 9D: SE-W, n=10; SE-J, n=7. 7W: SE-W, n=8; SE-J, n=8. CTRL epileptogenesis: CRTL-W, n=10; CRTL-J, n=6. CTRL epilepsy: CTRL-W, n=6; CTRL-J, n=6. Bonferroni *post-hoc* analysis following two-way ANOVA: ** *p*<0.01, *** *p*<0.001. Abbreviations: SE-W, SE induced at weaning; SE-J, SE induced at juvenile stage

**Fig. 6.**
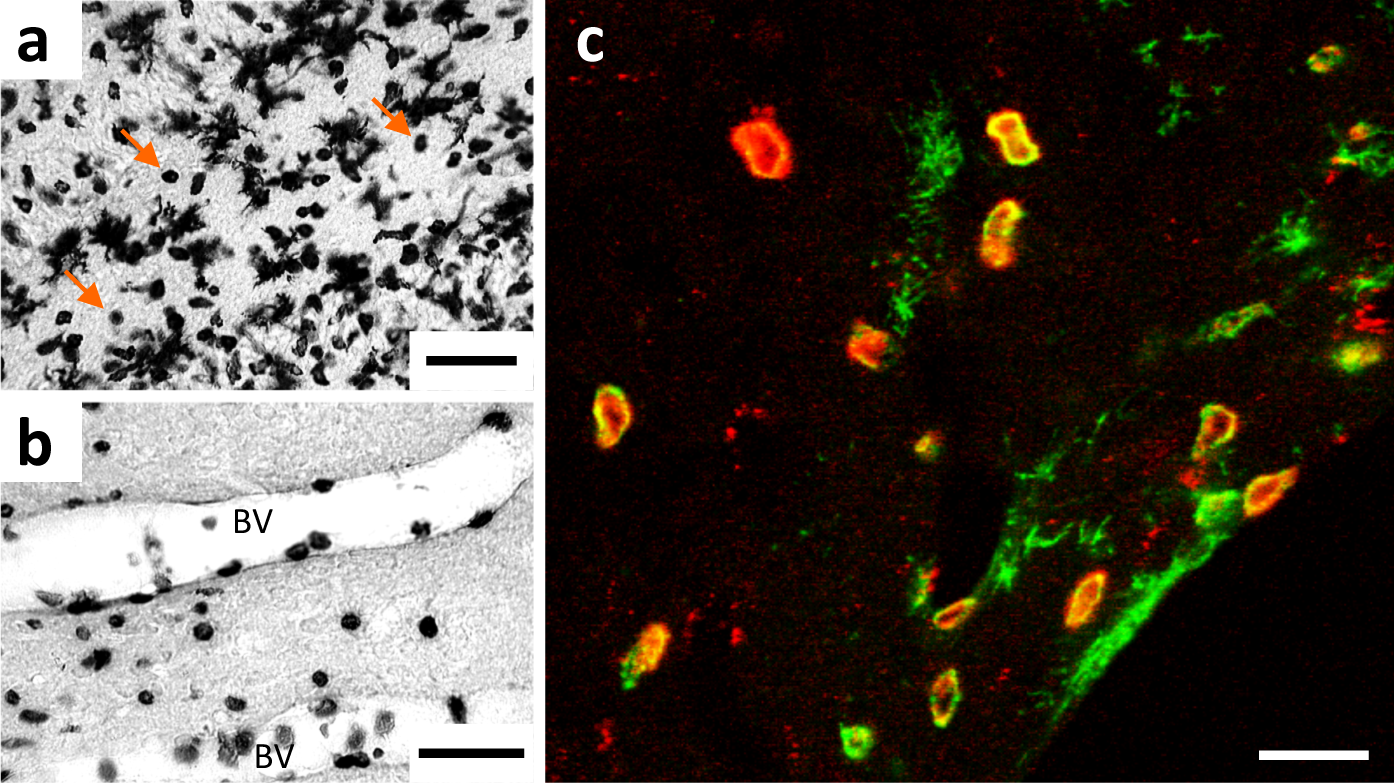
Immunohistological detection and identification of round-shaped cells expressing ITGAM (CD11b) 24 hours post-SE. (**a**,**b**) ITGAM was detected in the dentate gyrus of rats subjected to pilocarpine-induced SE at P42 and sacrificed 24 hours later. **a** Orange arrows depict “round-shaped cells” within the brain parenchyma, intermingled to activated resident microglial cells. **b** Firm adhesion of round-shaped cells to endothelial cells is illustrated. **c** Double fluorescent immunolabeling of ITGAM (Green) and CD14 (Red) in the hippocampus shows that almost all ITGAM+ cell infiltrates are monocytes/macrophages (CD14+). Scale bars: a,b: 50 µm; c: 20 µm

### Modeling of mTLE in rats suggests that some patients may have basal inflammatory levels in the hippocampus

As highlighted above, no control hippocampal tissues collected under similar conditions to those of mTLE patients were available to compare levels of inflammation measured in the resected hippocampi of mTLE patients to reference / baseline values. In this context, animal models of mTLE presented above have provided all their added value in that epileptic rats can be compared to control rats for which samples were obtained under perfectly identical conditions and, in addition, very similar to those of surgically resected tissues of mTLE patients, i.e. with quasi immediate freezing after tissue collection.

Transcripts level of the same panel of inflammatory mediators that were studied in mTLE patients were quantified in the hippocampus of SD rats that developed epilepsy after SE induced at weaning (EPI-W) or at juvenile stage (EPI-J) Measured values were compared to that of control rat groups (CTRL) (Fig. 7). The inflammatory levels of control rats (sacrificed at the same time as the epileptic animals, i.e. 7 weeks post-SE) whose SE was induced at weaning (CTRL-W) and juvenile (CTRL-J) stages were not statistically different, hence the two control groups were pooled in a same control group (CTRL). IL6 was not detected in any of the control rat samples.

**Fig. 7.**
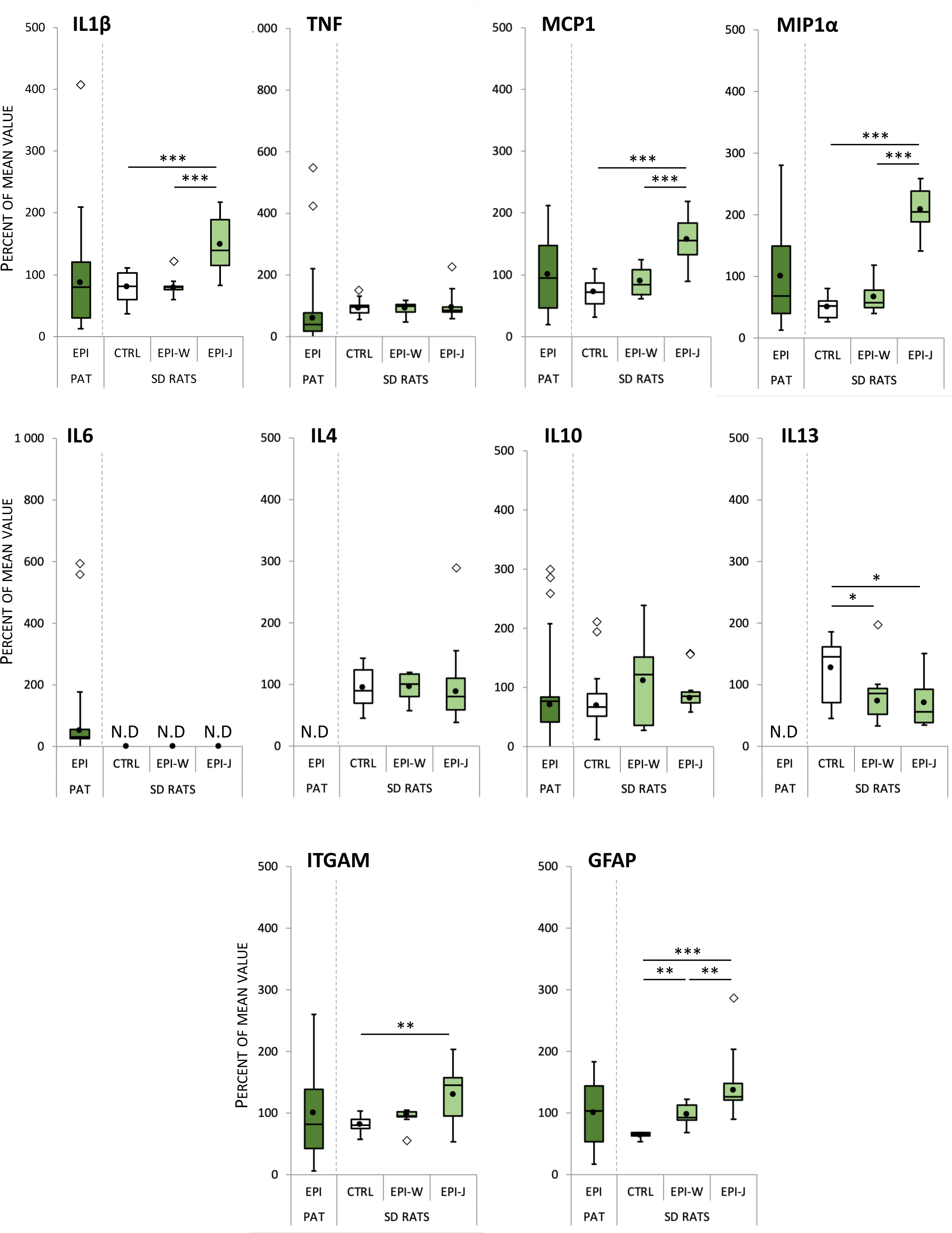
Heterogeneous distribution of inflammatory markers in the hippocampus of TLE patients can be modeled by the combination of two complementary models of TLE in rats. Distribution of the values of pro-inflammatory cytokines (IL1β, TNF, IL6), chemokines (MCP1, MIP1⍺), anti-inflammatory cytokines (IL4, IL10, IL13), and cellular markers (ITGAM, GFAP) in TLE patients (EPI-PAT) as well as in Sprague Dawley (SD) rats at the epileptic stage (7 weeks post-SE) following Pilo-SE induced at weaning (EPI-W, n=8) or at the juvenile stage (EPI-J, n=8) and in control SD rats (CTRL, n=12). Box-and-whisker plots model the distribution of each value around the median. Mean is represented by black dots. Outliers are represented by diamonds. Tukey’s *post-hoc* analysis following one-way ANOVA: * *p*<0.05, ** *p*<0.01, *** *p*<0.001. Abbreviations: N.D., not detected

In EPI-W rats, statistical analyses revealed that, except for IL13 (p=0.0384) and GFAP (p=0.0043), there was no significant difference between the dispersion of the CTRL group and the EPI-W group, indicating that when the SE is induced in weaned rats, the inflammation does not differ substantially from healthy rats. In contrast, in EPI-J, we show a significant difference between CTRL group and EPI-J group for IL1β (p=0.0002), MCP1 (p<0.0001), MIP1α (p<0.0001), IL13 (p=0.0234), ITGAM (p=0.0039) and GFAP (p<0.0001) (Fig. 7). We also demonstrate that EPI-J group is significantly different from EPI-W group for IL1β (p=0.0005), MCP1 (p=0.0005), MIP1α (p<0.0001) and GFAP (p=0.0034) (Fig. 7). No differences in expression of TNF, IL4 and IL10 were found between epileptic groups and control group as well as within epileptic groups.

In these two rat models of mTLE, we further investigated *il1β* gene regulation at protein levels and representative genes of the IL1 system, which is one of the most studied in the context of neuroinflammation in epilepsy. We tested the hypothesis of whether low levels of IL1β transcript in EPI-J rats could be due to a greater cytoplasmic pool of the corresponding protein that may exert a negative control on the transcription of this specific gene. We detected IL1β protein by immunohistofluorescence 7 weeks post-SE in controls and epileptic rats in both rat models (Fig. S5A-B). We found that IL1β protein was indisputably detected in EPI-J rats, whose IL1β transcript levels were significantly stronger than that of controls and of EPI-W rats (Fig. 7). By contrast, IL1β protein was barely detected in EPI-W rats, ruling out the hypothesis that low levels of IL1β-mRNA levels in these rats might be due to elevated cytoplasmic levels of IL1β protein. IL1β signaling depends on its target, interleukin-1 receptor type 1 (IL1R1) and its naturally occurring competitive IL1β receptor antagonist (IL1RA) [48]. Therefore, we measure transcript levels of these two genes, and found that both IL1R1 (Fig. S5C) and IL1RA (Fig. S5D) mRNA levels were induced in EPI-J rats compared to controls, but not in EPI-W rats. All these data support the hypothesis that key representative genes of the interleukin 1 system are similarly regulated. Although the numerous post-transcriptional mechanisms involved in the transformation of messenger RNAs into proteins are not yet sufficiently well defined to be able to predict protein concentrations from mRNA levels, our results indicate on the one hand that there is a coordinated expression of IL1β mRNA and protein, and, on the other hand, that genes representative of the interleukin 1 system appear to be less expressed in EPI-W rats than in EPI-J rats.

When considering both PI-I and IC-I, a strong difference was confirmed between the two models of epilepsy (Fig. 8), highlighting a significant difference between controls and EPI-W rats for the inflammatory cell index but not for the pro-inflammatory index. On average, the PI-I and the IC-I increased at most 1.96 times and 1.90 times, respectively, in epileptic rats compared to control rats (Fig. 8). Altogether, our preclinical data indicate that depending on the epilepsy model used, epilepsy can be associated or not with an induction of pro-inflammatory cytokines and chemokines (Fig. 8A), but is constantly associated with an induction of inflammation cell index (Fig. 8B) resulting from an induction of astroglial GFAP (Fig. 7).

**Fig. 8.**
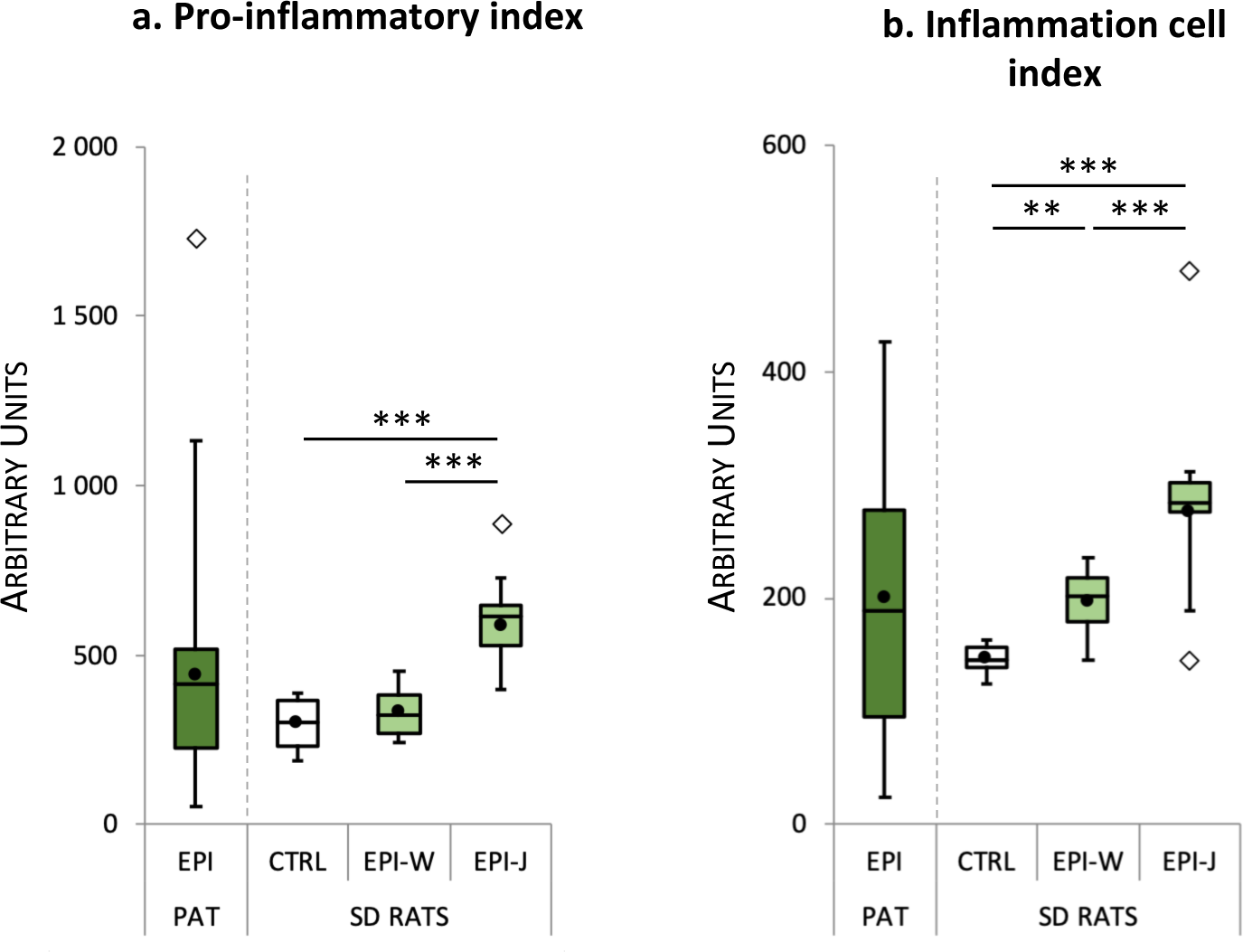
Indexes of inflammation in resected hippocampus of TLE patients and in epileptic rats. Pro-inflammatory **(a)** and inflammation cell **(b)** indexes in TLE patients (EPI-PAT) as well as in Sprague Dawley (SD) rats at the epileptic stage (7 weeks post-SE) following Pilo-SE induced at weaning (EPI-W, n=8) or at the juvenile stage (EPI-J, n=8) and in control SD rats (CTRL, n=12). Indexes were calculated from transcript levels as described in the methods section. Results are expressed as in Fig. 7. Tukey’s *post-hoc* analysis following one-way ANOVA: ** *p*<0.01, *** *p*<0.001

In the absence of reference values for samples from mTLE patients, we undertook a translational approach by facing off the data obtained in rats with those of patients, based on the dispersion of normalized values obtained for the different inflammatory markers. The total variability observed in rats, including both the control group and the two epileptic rat groups, covered between 49% and 93% of the variability observed in patients, with: 76% for IL1β, 49% for TNF, 93% for MCP1, 87% for MIP1α, 87% for IL10, 59% for ITGAM, 69% for GFAP (Fig. 7), 50% for the PI-I and 47% for the IC-I (Fig. 8). In addition, for each of the transcripts, the lowest normalized values were always observed in patients, and thus lower than the lowest values measured in control rats.

### Substantiate level of inflammation in the amygdala may counterbalance very low level of inflammation in the hippocampus of some mTLE patients

Since the 22 patients included in this study had undergone cortico-amygdalo-hippocampectomy, we also measured the pro-inflammatory index (Fig. 9) and the housekeeping gene index (Fig. S2) in the amygdala. As for the hippocampus (Fig. S2A), the housekeeping gene index is highly variable between patients within the amygdala (Fig. S2B), and is not consistent between the two brain structures (Fig. S2C). We show that the expression of pro-inflammatory genes in the amygdala is highly variable from one patient to another (Fig. 9A), and very high in some patients (e.g. P10 and P40; Fig. 9A). Interestingly, a comparison for each patient of the values measured in the hippocampus and the amygdala indicates that for some, the inflammation measured in the amygdala is stronger than that measured in the hippocampus (P40, x9.4; P41, x6.05; P16: x5.43; P17: 4.31; P1, x3.37). As a result, for some patients, the mean value measured in the amygdala-hippocampus complex appears higher than when only the hippocampus is considered, as in patient P40 (Fig. 9A). Finally, when we look at the inter-individual variations in the amygdalo-hippocampus pro-inflammatory index, they are much lower than when only the hippocampus or the amygdala is considered (Fig. 9B). The lowest amygdalo-hippocampus values were found to be always greater than the lowest values measured either in the hippocampus or the amygdala (Fig. 9). Finally, as for the hippocampus, we found that using housekeeping genes instead of the SmRNA to normalize the RT reaction would have led to biased results (Fig. S3B).

**Fig. 9.**
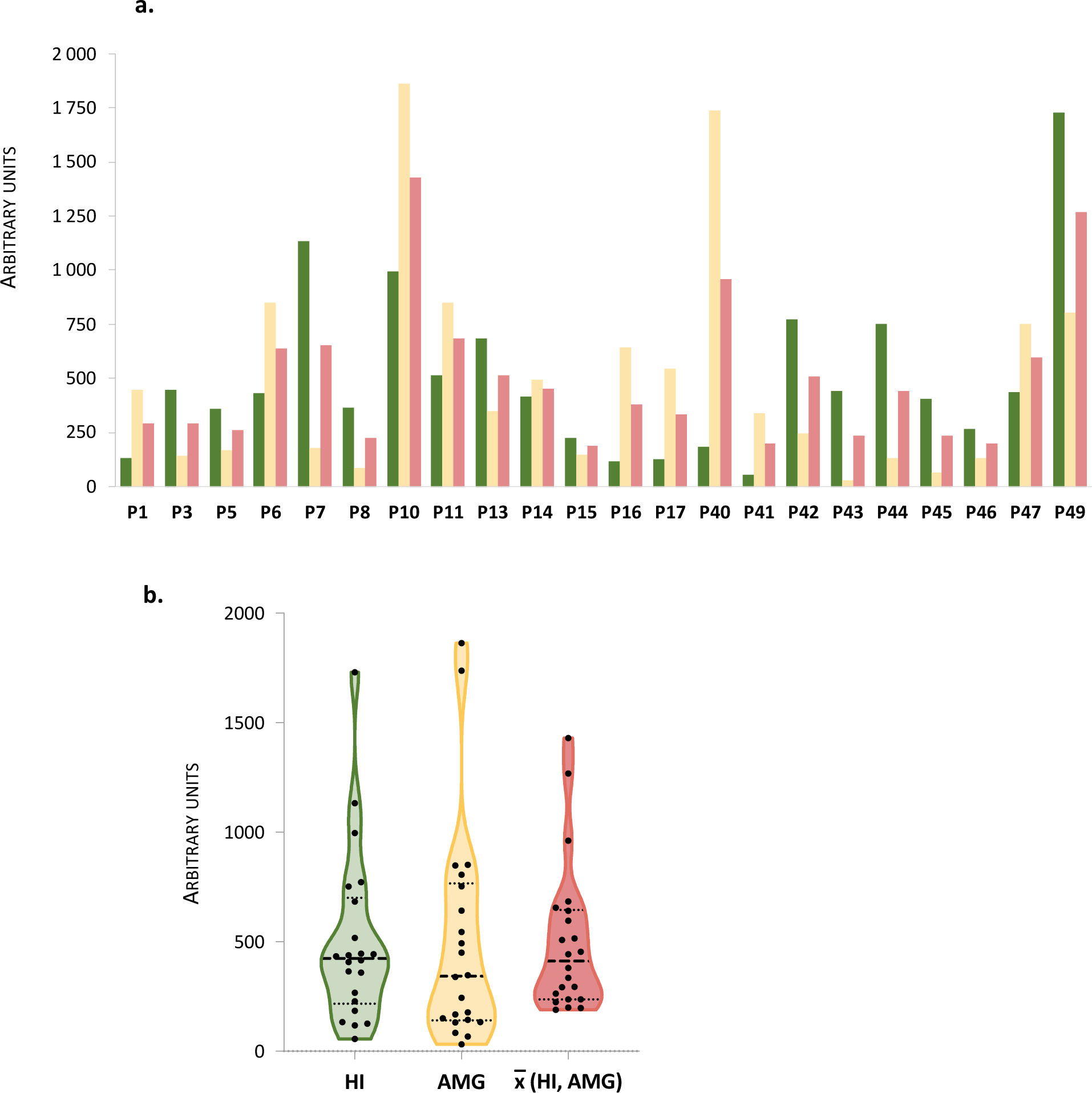
Pro-inflammatory index in resected amygdala of TLE patients compared with the hippocampus. **a** The pro-inflammatory indexes measured in the hippocampus (green bars) and the amygdala (yellow bars) have been averaged (red bars) to evaluate the inflammatory status in the whole amygdalo-hippocampal complex. **b** Violin plot displaying the pro-inflammatory index distribution of the data illustrated in **(a)**. Distribution of each value (black dots) is plotted around the median (dashed line) and the 25 and 75 percentiles (dotted lines)

The amygdalo-hippocampus pro-inflammatory index (PI-I, in A.U.) did not correlate either with the age (in year) at epilepsy onset (PI-I = 6.33 x age + 433.2; r^2^=0.0285) nor with the duration (in year) of epilepsy (PI-I = 1.364 x duration + 472.7; r^2^ = 0.0021).

### Inflammation is of low grade in chronic epilepsy compared to explosive inflammation during epileptogenesis

The ∼ two-fold increase in the pro-inflammatory index in rats with epilepsy developed after SE induced at the juvenile age (EPI-J) (Fig. 8A) raised the issue of whether this increase was greater or lesser than that occurring after SE itself, as already questioned using undisputable quantitative procedures following kainic acid-induced SE [18]. To this end, inflammatory levels in our two models of mTLE were investigated during epileptogenesis and compared to those measured during chronic epilepsy. Transcript levels of inflammatory and anti-inflammatory markers were quantified in the hippocampus of rats during epileptogenesis (at 7 hours, 1 day, 9 days) and during epilepsy (7 weeks) after the onset of SE induced at P21 (SE-W, light blue bars) or P42 (SE-J, dark blue bars). The results are presented for each pro-inflammatory (Fig. 10) and anti-inflammatory (Fig. 11) markers, as well as for the corresponding pro-inflammatory and anti-inflammatory indexes (Fig. 12). They reveal that the induction peak occurred between 7 hours and 1 day after SE for both epileptic models. The comparison of the peak values of pro-inflammatory markers determined during epileptogenesis with the values measured during the chronic phase of epilepsy (7 weeks post-SE) reveals that the difference between these two values ranged between 0.46-fold (TNF) and 740-fold (MCP1) for rats subjected to SE at weaning (P21) and between 8-fold (TNF) and 781-fold (MCP1) for rats subjected to SE at the juvenile stage (P42) (Table 4). Hence, the pro-inflammatory index measured at the peak during epileptogenesis was 17.77- and 23.95-fold greater than that measured in the chronic phase of epilepsy, for SE induced at P21 and P42, respectively (Table 4).

**Fig. 10.**
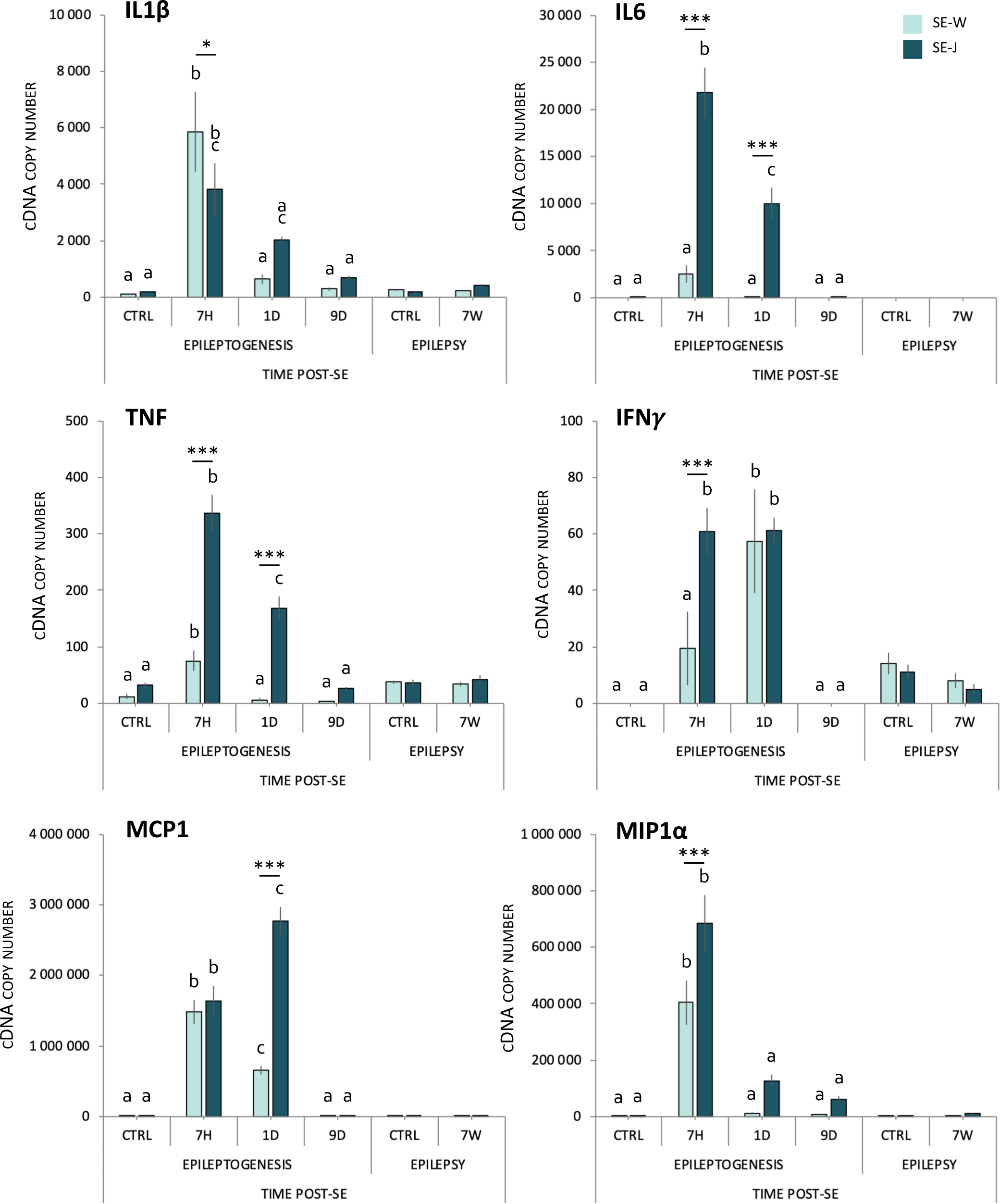
Transcript levels of pro-inflammatory cytokines and chemokines after Pilo-SE. Transcript values of pro-inflammatory cytokines (IL1β, TNF, IL6, IFN*ɣ*) and chemokines (MCP1, MIP1⍺), during epileptogenesis, i.e at 7 hours (7H), 1 day (1D), 9 days (9D) post-SE and once epilepsy was chronically installed, i.e. 7 weeks post-SE (7W) compared to respective controls. Results are expressed as in Fig. 5. Bonferroni *post-hoc* analysis following two-way ANOVA: * *p*<0.05, *** *p*<0.001. Abbreviations: SE-W, SE induced at weaning; SE-J, SE in induced at juvenile stage

**Fig. 11.**
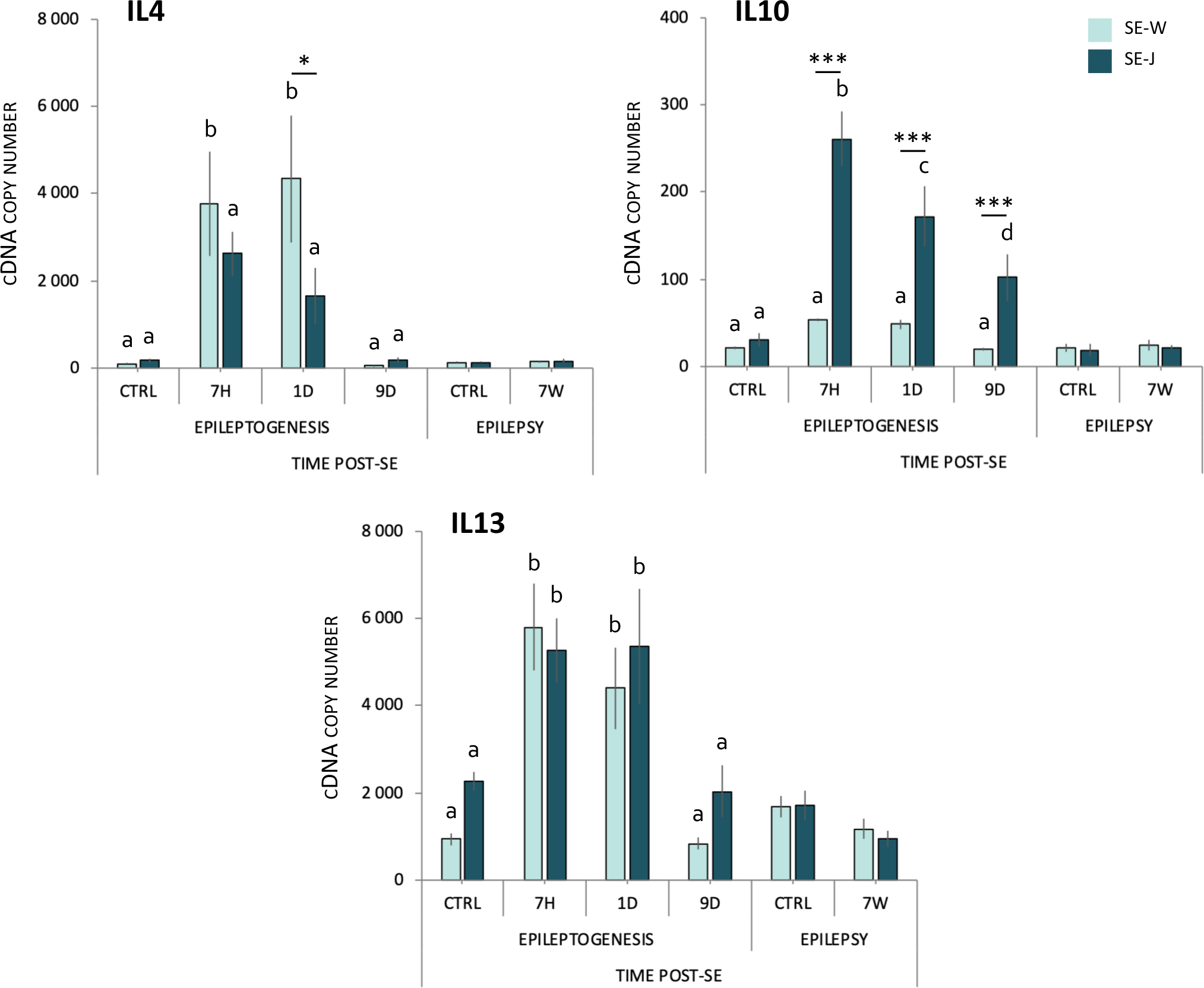
Transcript levels of anti-inflammatory cytokines after pilocarpine-induced SE. Transcript values of anti-inflammatory cytokines (IL4, IL10, IL13), during epileptogenesis, i.e at 7 hours (7H), 1 day (1D), 9 days (9D) post-SE and once epilepsy was chronically installed, i.e. 7 weeks post-SE (7W) compared to respective controls. Results are expressed as in Fig. 5. Bonferroni *post-hoc* analysis following two-way ANOVA: * *p*<0.05, *** *p*<0.001. Abbreviations: SE-W, SE induced at weaning; SE-J, SE in induced at juvenile stage

**Fig. 12.**
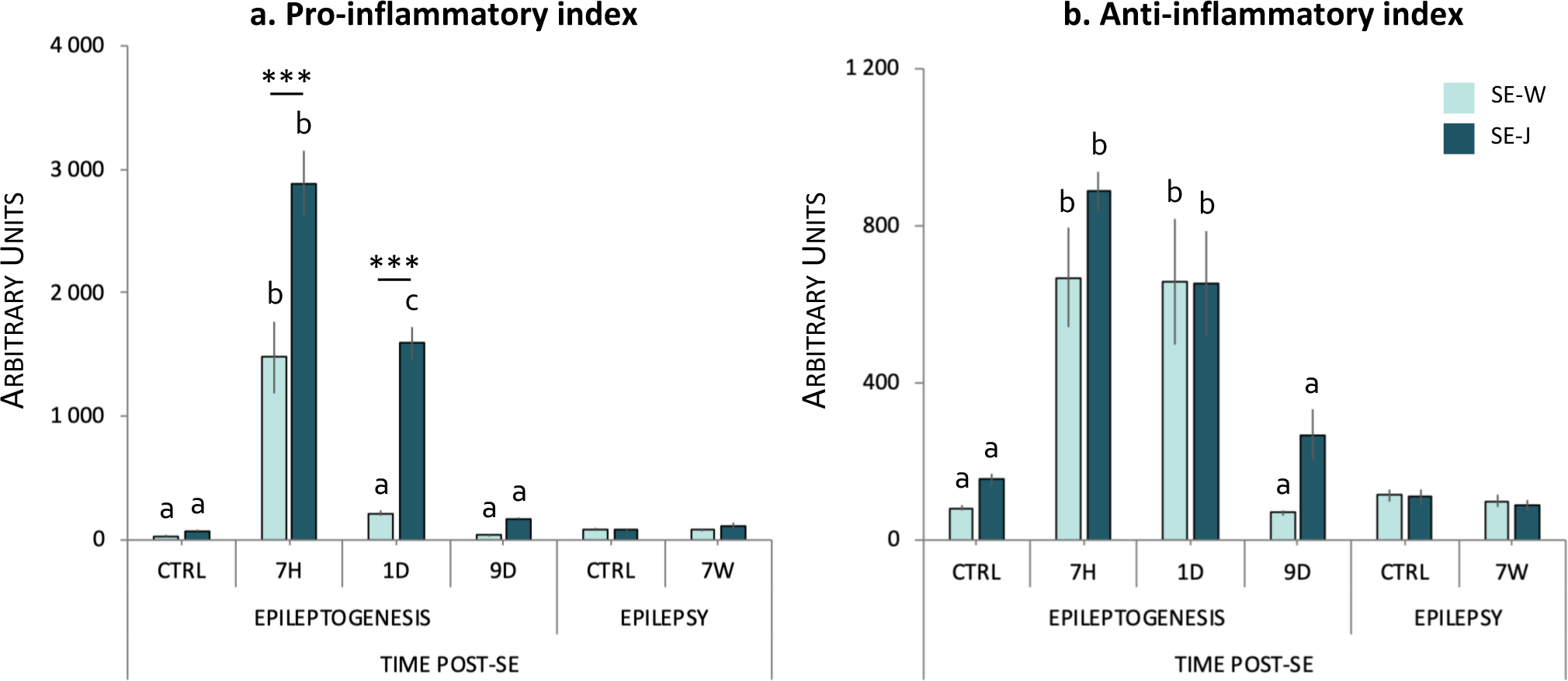
Inflammation during epilepsy is of low-grade compared to that during epileptogenesis. Pro-inflammatory **(a)** and anti-inflammatory **(b)** indexes in Sprague-Dawley (SD) rats were calculated during epileptogenesis, i.e at 7 hours (7H), 1 day (1D), 9 days (9D) post-SE and once epilepsy was chronically installed, i.e. 7 weeks post-SE (7W) compared to respective controls. Results are expressed as in Fig. 5. Bonferroni *post-hoc* analysis following two-way ANOVA: *** *p*<0.001. Abbreviations: SE-W, SE induced at weaning; SE-J, SE in induced at juvenile stage

**Table 4.**
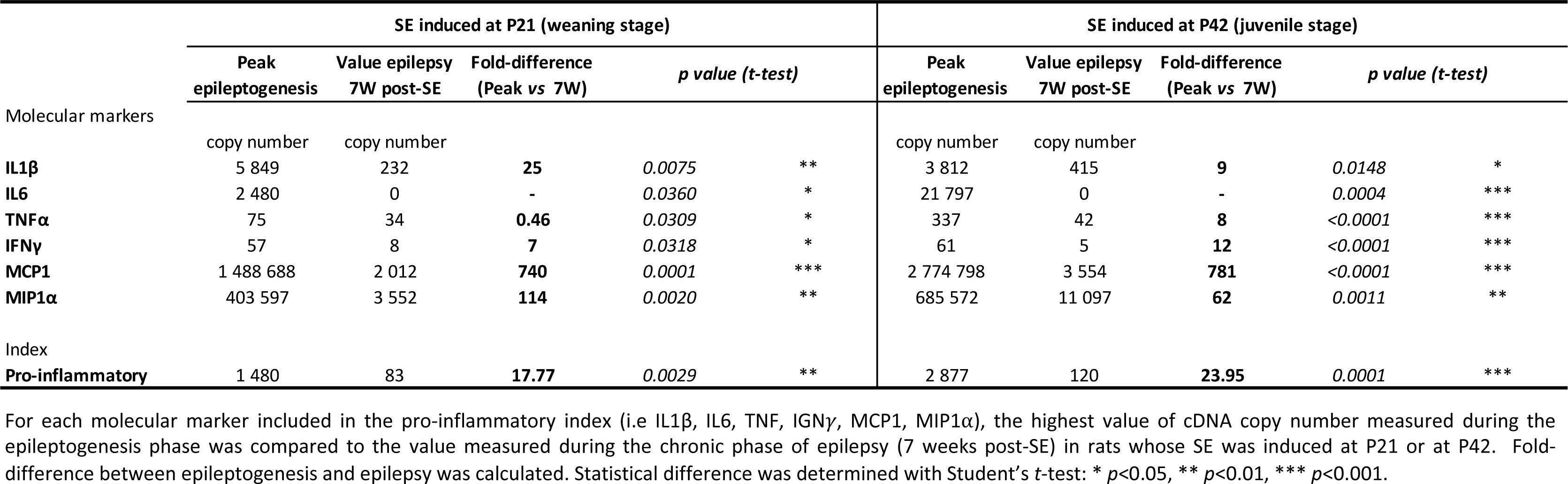
Fold-changes in inflammatory markers between epileptogenesis and epilepsy in rats. For each molecular marker included in the pro-inflammatory index (i.e IL1β, IL6, TNF⍺, IGN*ɣ*, MCP1, MIP1⍺), the highest value of cDNA copy number measured during the epileptogenesis phase was compared to the value measured during the chronic phase of epilepsy (7 weeks post-SE) in rats whose SE was induced at P21 or at P42. Fold difference between epileptogenesis and epilepsy was calculated. Statistical difference was determined with Student’s *t-*test: * *p*<0.05, ** *p*<0.01, *** *p*<0.001

### Quantitative RNAscope® *in situ* hybridization confirms data obtained by RT-qPCR

Data on transcript levels acquired so far in this study were obtained by RT-qPCR. They indicate wide variations for most of the studied inflammation markers in the hippocampus both between patients and between different groups of rats, especially for the latter between the epileptogenesis period and the chronic phase of epilepsy. In order to rule out any hypothesis that the observed variations could be the result of random degradation of mRNAs during the extraction and purification phases of total RNAs, RT-qPCR data for IL1β were compared to those obtained on fixed brain sections by the RNAscope® technology, which is a highly quantitative *in situ* hybridization (ISH) method. IL1β was selected for this comparison because it is one of the most studied inflammation markers in the context of neuroinflammation. Finally, to carry out this comparison, we selected 5 patients whose IL1β-cDNA copy numbers were either low (P15 and P45), intermediate (P43 and P44) or high (P42), and the different time points studied by RT-qPCR during epileptogenesis and the chronic phase of epilepsy in rats subjected to Pilo-SE at the juvenile (P42) stage. In sections of the hippocampus resected from mTLE patients, the density of IL1β-mRNA signal (magenta dots) was greater in patient P42 compared to patient P15 (Fig. S6), as expected, and the surface area occupied by IL1β-mRNA signal in the 5 selected patients correlated significantly with the corresponding IL1β-cDNA copy numbers quantified by RT-qPCR (Fig. 13A). In rats subjected to SE at P42, IL1β-mRNA signal was quantified in the dentate gyrus where it was greater than other regions of the hippocampus, including CA1 area. The density of IL1β-mRNA signal, measured at 7 hours, 1 day, 9 days and 7 weeks post-SE, was greater at 7 hours post-SE (Fig. S7 and Fig. 13B1) and significantly correlated with IL1β-cDNA copy numbers quantified by RT-qPCR in different sets of animals (Fig. 13B2).

**Fig. 13.**
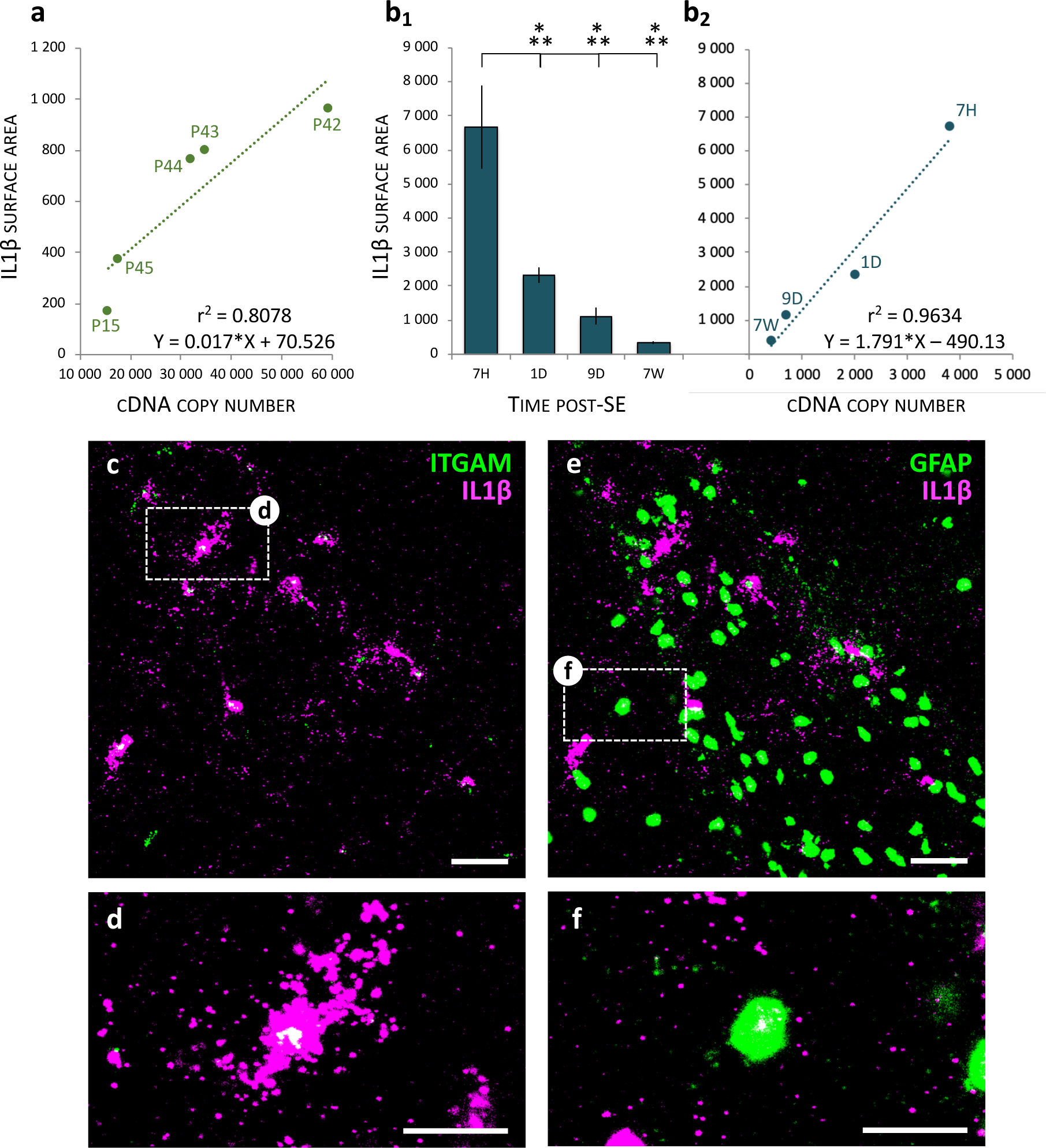
RNAscope® ISH of IL1β-mRNA confirms RT-qPCR data and reveals in rats subjected to Pilo-SE that IL1β-mRNA is mainly expressed by microglia at the peak of inflammation. **a** Scatter plot between IL1β cDNA copy number measured by RT-qPCR in the hippocampus of TLE patients and the surface area occupied by IL1β-transcript signal in sections processed by RNAscope® ISH. Data are obtained from the 5 patients, whose resected hippocampi were split in two parts, one reserved for RT-qPCR, the other one for histology. Data are significantly correlated and fitted by a linear regression, p<0.0381. **b** Quantitation of the surface area occupied by IL1β-transcript signal in the granule cell layer of the dentate gyrus of rat brain sections processed by RNAscope® ISH **(b1)**. Sections were selected at Bregma −4.16 mm from rats sacrificed during epileptogenesis (7 hours (7H), 1 day (1D), 9D after SE) or during chronic epilepsy (7 weeks (7W) after SE). Statistical analyses showed significant differences between the IL1β surface area measured 7H post SE (n=4) and all the other time points (1D: n=5; 9D: n=3; 7W: n=5). Scatter plot **(b2)** between the average IL1β cDNA copy number determined by RT-qPCR in the hippocampus of rats sacrificed at the same time points as in B1 (Fig. 10) and the average surface area occupied by IL1β-transcript signal measured in sections processed by RNAscope® ISH **(b1)**. Data are significantly correlated and fitted by a linear regression, p<0.0194. **(c-f)** Triple *in situ* hybridization of IL1β together with ITGAM **(c-d)** and GFAP **(e-f)** transcripts using RNAscope® technology, in the dentate gyrus of the rat hippocampus 7 hours (peak of inflammation; Fig. 10) after SE induced at 42 days. To facilitate the visualization of IL1β in microglia and astrocytes, we used two colors providing the best contrasts, and thus assigned magenta to IL1β and green either to ITGAM (microglia) or GFAP (astrocytes). Colocation is displayed in white when magenta and green are superimposed. In this area, the largest amount of IL1β transcript is colocalized with ITGAM+ cells **(c-d)**, compared to astrocytes **(e-f)**. Confocal microscope images are magnified at 63X. Scale bars: C and E: 50 *µ*m; D and F: 25 *µ*m

### Microglial cells seem to produce most of the IL1β during the acute phase

Previous studies using immunohistochemical procedures reported that IL1β was expressed mainly by astrocytes and more rarely by microglial cells/macrophages in the hippocampus following pilocarpine-induced SE and self-sustained limbic SE [36], and exclusively by astrocytes following kainic acid-induced SE [18]. To determine to which extent microglial cells or astrocytes were each involved in the production of IL1β-mRNA, we used multiplex detection of IL1β, ITGAM and GFAP transcripts using RNAscope® ISH. We could not combine IL1β-mRNA ISH with immunohistofluorescent detection of GFAP and ITGAM because antigens recognized by the different antibodies tested were altered by the permeabilization and fixation procedures in RNAscope® protocols. In patient P42, who had the greatest IL1β-cDNA copy number, IL1β-mRNA signal (magenta dots) was located in cells bearing morphological features of glial cells (Fig. S6A). However, the paucity of ITGAM-mRNA and GFAP-mRNA signals at the location of IL1β-mRNA signal precluded in mTLE patients the identification of IL1β-mRNA signal as being of astroglial or microglial origin (Fig. S6B-C). In rats, the only time point it was possible to identify the glial cells expressing IL1β-mRNA was 7 hours post-SE. Cells with a large and packed IL1β-mRNA signal appeared to be ramified microglial cells, as identified by the presence of ITGAM-mRNA signal in the core of the IL1β-mRNA signal (Fig. 13C-D). At this time point, numerous astrocytes also expressed IL1β-mRNA, but at weaker levels compared to ITGAM+ cells, as depicted by the small surface area occupied by IL1β-mRNA signal with the dense signal corresponding to GFAP-mRNA (Fig. 13E-F).

## 4 Discussion

The current study reports that inflammation in resected hippocampus of patients with mTLE presented with a high inter-individual variability. Such a variability was also found in the amygdala. Our intriguing result showing that short-term delay of resected tissue processing led to large decrease of housekeeping gene mRNAs, precluded the possibility of using post-mortem tissues to estimate physiological baseline transcript levels in non-epileptic tissues, and then to evaluate the degree of the inflammatory status in the resected tissues of mTLE patients. To overcome this problem, as an alternative, we used mTLE models in rats developing epilepsy after a SE induced either at weaning or at the juvenile stage, These animal models have the advantage of providing access to healthy control brain tissues. We show that the data obtained in epileptic rats model a large part of the variability observed in patients. In addition, in the chronic phase of epilepsy, the levels of selected neuroinflammatory markers measured in the hippocampus varied between values ranging from 0.87 to 9.55 times those of controls. We also showed that inflammation during the chronic phase, when present, is of low grade compared to that measured after an epileptogenic brain insult. Finally, we demonstrated that microglial cells are the main contributors to the production of interleukin-1β during the acute phase after SE.

### Methodological considerations

In our study, we chose to evaluate gene expression at the transcript level rather than at the protein level, because the method mostly used to quantify RNA (calibrated RT and real-time PCR) is much more quantitative than those used for proteins quantification (Western Blot and Elisa). Indeed, the two most common methods for protein quantification depend on the availability of validated antibodies for each of the targeted genes. In addition, we showed in this study that a given protein (GFAP) did not show the same tissue distribution pattern when detected with two distinct specific antibodies. By contrast, when considering RNAs, even if amplification of given cDNAs by PCR requires different primer pairs between humans and rats, it remains highly specific to the corresponding mRNA. Furthermore, PCR is quantitative as soon as it is performed on a real-time thermocycler and a calibration curve is used, giving access to the number of cDNA copies detected. To generate the cDNAs to be amplified by PCR, we have chosen a method that allows us to calibrate reverse transcription using a synthetic and exogenous poly-A RNA (SmRNA) (WO20040404092414) [13, 29, 30, 39]. This contrasts with the selection of one or more endogenous genes, so-called housekeeping genes, considered as internal controls and, *de facto*, as being *a priori* invariant in all studies that use this kind of standardization. Our methodological approach is all the more justified when considering our results showing that three mostly used housekeeping genes greatly vary between patients.

When several mRNAs of inflammation markers are quantified, and some of them vary upward while others vary downward, one of the major difficulties is to define whether the overall level of inflammation has increased or decreased. For this reason, inspired by Gene Ontology analysis, we have established three indexes to report “global” pro-inflammatory, anti-inflammatory and glial activation states, based on a number of mRNAs for which we also provided individual quantifications. To generate theses indexes, we have chosen prototypical pro-inflammatory cytokines and chemokines IL1β, IL6, TNF, MCP1 and MIP1α involved in epilepsy pathophysiology [3, 6, 48], anti-inflammatory cytokines IL4, IL10 and IL13 which expression is increased in various neurological pathologies [19, 26, 28], and finally GFAP and ITGAM, which are respective markers of astrogliosis and microgliosis, both involved in epilepsy [11].

### Post-mortem tissues

Inflammation has for years been considered as a key contributor to the pathophysiology of epilepsy [46], which encouraged several studies to investigate the degree of neuroinflammation in the epileptic brain. One of the commonalities between most of these studies, regardless of the quantification methodology employed, has been the use of post-mortem tissue obtained from autopsy non-epileptic control subjects to compare with values measured in specimen of epileptic patients. Although not epileptic, some individuals suffered from other neurological conditions such as brain tumor or had experienced traumatic injuries, raising concerns about the inflammatory status of these samples in comparison with healthy tissues [15, 41]. Another issue is the delay of processing of these post-mortem tissues, ranging from 4 to 20.5 hours [2, 10, 15, 21, 23, 31, 36, 40, 41]. While earlier studies have shown that RNA can remain substantially intact, even for long periods of time after death [16, 20, 34], other studies have reported that post-mortem interval should be controlled in human and animal models [5], especially for mRNA profiling studies [14, 45]. In addition, a recent study has provided evidence that data obtained for miRNAs extracted from resected tissues of epileptic patients were different to those of post-mortem tissues from epileptic patients [37]. A further concern not raised so far about the use of post-mortem human tissues in general is related to the heterogeneity of the individuals included in a study. Thus, adding over to this heterogeneity the variability related to the uncontrolled degradation of the mRNAs only adds uncertainty to the data produced.

Our results obtained on hippocampal tissues from 10 patients with epilepsy showed that a short (45-90 min) delay in the processing of a sample, although handled according to standard procedures, resulted in substantial decrease in three housekeeping gene transcript levels. This may reflect a specific degradation of mRNAs, not detected by the widely used reference method based on the integrity of two very abundant ribosomal RNAs. For all these above-mentioned reasons, we preferred not to use post-mortem tissues to define basal inflammatory levels in the hippocampus.

### Variable levels of inflammation in the hippocampus of mTLE patients

While several studies have reported increased inflammation in the resected hippocampus of mTLE patients when compared to post-mortem controls [15, 21, 31, 36, 52], greater IL1β and IL6 levels have also been measured in the resected hippocampus of non-epileptic patients compared to mTLE patients [41]. These conflicting data fuel the debate about whether substantial inflammation is present in the epileptic focus [2]. In our patient study, we do not have physiological baseline mRNA levels of the targeted cytokines and chemokines due to the lack of appropriate human control tissues. However, we show that the variations of the mRNA levels were highly variable in resected hippocampi of mTLE patients, certain markers of inflammation being undetectable in some while they were very easily detectable in others. Since it has been shown that brain structures involved in the epileptic network, but at a distance from the epileptic focus, may have higher levels of inflammation than in the epileptic focus itself [41], we also measured inflammation levels in the amygdala of the 22 patients included in our study. Inter-individual variations in the pro-inflammatory and cell inflammation indexes have also been found in the amygdala but without correlation between the hippocampus and the amygdala. A high level of inflammation in the amygdala may thus be associated with a low level in the hippocampus, or vice versa. This is particularly the case for two patients who had radically opposite levels of inflammatory marker expression between the two brain structures (e.g. Patient 7 and Patient 40). These variations did not correlate with gender, age, duration of epilepsy, and ASDs. Our study is thus in support of the hypothesis that inflammation should be investigated not only at the site of the epileptic hippocampus, but also within the entire epileptic network to have a more integrative overview of the brain inflammatory status in mTLE.

An important question is whether there is a strong link between inflammation in the hippocampus and the recurrence of seizures. In our study, this question could not be answered in rats, since our preliminary studies showed that the sole implantation of screws into the skull induced long lasting brain inflammation in the underneath cortex, as well as in the hippocampus (data not shown). In the human part of our study, although data on seizure frequency have only been obtained in 5 patients, they do not indicate a positive correlation between pro-inflammatory cytokine levels in the hippocampus and seizure frequency. A PET study evaluating microglial activation in a mTLE patient showed that it was greater 36 hours after the last seizure compared to a seizure-free period [4]. It cannot thus be excluded that patients with the highest inflammatory levels are those who experienced the most recent seizures.

Our choice to model mTLE in rats after SE induced by pilocarpine provided us the possibility to have access to physiological baseline values for the different markers of inflammation measured in the hippocampus. When taking the pro-inflammatory and anti-inflammatory indexes, or the individual mRNAs constituting these indexes, our results show that once epilepsy was developed, rats whose SE was triggered at weaning (P21) were not distinguishable from controls, unlike rats whose SE was induced at the juvenile stage (P42). Since some epileptic rats had similar levels of inflammation to that of control rats, we established the inter-indivudual variability that exists in the entire cohort of rats, including both controls and all epileptic rats. Given the translational approach of our study, we compared this inter-individual variability to that established between the 22 patients with mTLE. Intriguingly, the inter-individual variability in the rats almost overlapped the variability observed between patients with mTLE. This comparison suggests that epilepsy may be active despite a barely detectable level of inflammation, at least as represented by the selected genes.

### Inflammation in epilepsy is low grade compared to that measured in the acute phase after status epilepticus

One of the major added values of our study is to have been able to quantify with the exact same methodology the mRNAs of some prototypical markers of inflammation, both in mTLE patients and in rats in different phases of epileptogenisis following pilocarpine-induced SE. In addition to providing us with valuable controls to establish physiological baseline levels of inflammation, the advantage of the animal model is to give us access to the entire period of epileptogenesis following SE, which is of course impossible in humans.

Very few studies have reported variations in mRNA levels of prototypic markers of inflammation during epileptogenesis up to epilepsy onset, with a method as quantitative as RT-qPCR. Recently, changes in IL1β and TNF mRNA levels were quantified in the hippocampus in a mouse model of mTLE developed after SE induced by intrahippocampal administration of kainic acid. Although the onset of epilepsy in this model was rapid (<7 days), quantification was limited to the first week post-SE and showed that the apparent peak of inflammation was between 2h and 72h, with values at 7 days still very high but not statistically different from that of controls [18]. Maximum increases, corrected by 3 reference genes, were in the range of 15 to 50-fold compared to controls for IL1β and TNF, respectively. In our study, after induction of SE by pilocarpine, the maximum increases reported, without the use of reference genes thanks to the use of an external calibrator, were of an order of magnitude equivalent to that of the intrahippocampal kainic acid (KA) model in mice, ranging from 6 to 55-fold relative to controls for TNF and IL1β, respectively. It is thus clear that the degree of inflammation in the hippocampus of epileptic rats, 7 weeks after SE induced at P42, was of low-grade compared to the explosive inflammation measured in the first hours to days following SE.

### The extent of neurodegeneration and gliosis might depend on the degree of inflammation in response to an epileptogenic brain insult and the delay to recover baseline levels

Prior studies have shown that the inflammation in the hippocampus of mTLE patients was not greater in the presence of hippocampal sclerosis [2, 21]. Here, we provide evidence that patients who had the highest values of pro-inflammatory index (P07, P10, P13, P44 and P49) were those with the greatest scores of neuronal loss and reactive gliosis. In our animal study, only epileptic rats whose SE was induced at P42 had a pro-inflammatory index above that of controls and massive lesions in cortico-limbic and thalamic areas (P42) [8, 9, 29, 39, 51], compared to epileptic rats whose SE was induced at P21. The comparison of SE induced at P21 and P42 highlighted that the peak of the pro-inflammatory index was higher at P42 than at P21, with a slower return to baseline values, suggesting that a higher and longer exposure to inflammatory molecules might partly explain the massive neurodegenerative processes and gliosis observed in epileptic rats when SE was induced at P42. It is noteworthy that in epileptic rats subjected to pilocarpine-induced SE at P42, the apparent peaks were much more transient (between 7h and 24h post-SE) than those observed in the intrahippocampal KA mouse model that present with massive neurodegerative processes and gliosis [18]. This suggests that the extent of neurodegeneration and gliosis in mTLE models is dependent on the peak of inflammation and the time to return to baseline following the epileptogenic brain insult; this might also be the case in patients with mTLE.

### Which brain cells contribute the most to neuroinflammation?

Of the three cytokines studied (IL1β, IL6 and TNF), IL1β was the one that was still at a higher level than controls in rats that developed epilepsy after the induction of SE at P42. To identify which cells expressed IL1β, we had initially opted for double immunohistological labeling, but the radically different results obtained with the different anti-IL1β antibodies tested (none of which were validated) led us to go with quantitative RNAscope® in situ hybridization. In accordance with RT-qPCR results, maximum signal was observed 7h post-SE, and was clearly located in cells expressing Itgam-mRNA, with a morphology resembling that of activated microglial cells. Very small amounts of IL1β-mRNA were also detected in astrocytes at 7h post-SE. Due to the rapid decrease in IL1β-mRNA levels following SE, the dispersion of the corresponding signal did not make it possible to identify whether IL1β-mRNA was expressed in microglial cells or astrocytes. According to the results of previous studies, even if IL1β-immunopositive cells have been shown to resemble activated microglial cells in the hours following SE induced by intrahippocampal KA in rats [47], IL1β has been shown to be expressed in the hippocampus mainly by astrocytes at all phases of epileptogenesis and once epilepsy has developed in rats after pilocarpine-induced SE or after self-sustained SE [36] or at epilepsy onset in mice subjected to SE after intrahippocampal KA administration [18].

While studies in mice after pilocarpine-induced SE indicate that myeloid infiltrates (essentially macrophages) are responsible for the majority of the pro-inflammatory cytokines measured in brain tissue in the acute phase (24h-96h) post-SE [44, 50], our data acquired in rats subjected to pilocarpine-induced SE show that the inflammatory peak occurred 7h post-SE, at a time when no myeloid infiltrate was detected, whereas when these infiltrates were present between 24 and 48 hours post-SE, mRNA levels of pro-inflammatory cytokines were dramatically decreasing. In addition, the detection of IL1β by RNAscope® *in situ* hybridization did not make it possible to demonstrate, 24 hours post-SE, a stronger signal in round-shaped cells, resembling infiltrating macrophages. Therefore, if our *in situ* quantification methods are correct, one must consider that either the contribution of macrophages to brain inflammation following SE is radically different between rat and mouse models, or that the rather long procedures needed for separating microglial cells from macrophages by FACS in mice differently affected the turnover of cytokine mRNAs, leading to the differences observed between the two populations of cells.

### Translational relevance of the study

One of the major problems in studies involving brain biopsies taken from patients is that while the data obtained from these biopsies can be compared with each other, they cannot be compared with data that would have been obtained in an equivalent manner from healthy subjects. The results of our study clearly show that a delay of approximately 45-90 min between the surgical resection of the tissue and its freezing dramatically compromises the quantity of mRNAs. We have therefore chosen not to use hippocampi or brain tissues obtained post mortem, which mostly takes well over 90 min after death. Just as we decided not to use brain tissue from patients with pathologies other than epilepsy. With all the necessary precautions, in order to define what could be the level of inflammation in the resected tissues of patients with mTLE, in comparison to physiological baseline level on the one hand, and in comparison to the explosive inflammatory conditions generally observed in the acute phase of severe brain insults on the other hand, we have chosen to exploit to the maximum the predictive value that animal models can provide. To do so, we compared the inter-individual variability observed in the complete cohort of rats, including both control rats and two complementary models of epileptic rats, to that observed in patients with mTLE. This allowed us to propose that the level of inflammation in the hippocampus of patients with mTLE was low-grade, and that some may even have active epilepsy with inflammation levels almost identical to those of controls.

Our amygdala data also show that brain inflammation in mTLE must be looked at beyond the hippocampus, throughout the entire brain network involved in epilepsy. At this stage, the question that arises is how to measure the level of brain inflammation non-invasively, and, in the best case, from peripheral biomarkers [46].

### Limitations of the study

One of the major limitations of our study is that we were unable to measure the EEG activity of rats, by fear of inducing significant inflammatory levels at the hippocampal level, which could themselves have modified the course of the disease.

In our study, we explored inflammation by measuring the most studied prototypical cytokines and chemokines in epilepsy [43, 48]. However, the members of these classes of molecules extend far beyond those we have studied [52]. In addition, eicosonoids, which are metabolic derivatives of arachidonic acid, play a major role in inflammatory signaling and were not examined in this study, whereas their deregulation has clearly been identified in epilepsy [46].

Targeting mRNAs by RT-qPCR is certainly one of the most accessible methods of measuring gene expression in the most quantitative and reliable way possible. However, variations in the corresponding proteins would have certainly provided more relevant information on the most active signaling pathways in epileptic tissue. Easier access to high-throughput proteomics should help solve this issue, at least in part. It remains that we had trouble identifying the cells expressing cytokines at all stages of the development of the disease in animal models and on the resected hippocampi of mTLE patients. Such identification of cells expressing the proteins of interest will depend on the development of more specific and better validated antibodies.

## Supporting information

N.Gasmi_Supplementary figures and tables

## ACKNOWLEDGEMENTS

Nadia Gasmi was granted a PhD fellowship from the Fondation pour la Recherche Médicale (FRM grant number ECO20160736074 to NG).

## 10 Supplementary Material

**Fig. S1 Non-overlapping of GFAP immunofluorescent labelling obtained with two different antibodies.** Double immunohistochemical labelling of GFAP in the hippocampus (**a**) and in the piriform cortex (**b**) of a rat 7 weeks after pilocarpine-induced SE. The rabbit polyclonal anti-GFAP antibody (AB5804; Chemicon) is visualized in green while the mouse monoclonal anti-GFAP antibody (G3893; Sigma-Aldrich) is visualized in red. Colocalization is displayed in yellow when red and green are superimposed. These observations suggest that GFAP epitopes recognized by the two antibodies are not accessible (or present) in the same manner within the same structure or between two different structures. Scale bar: 50 µm

**Fig. S2 Index of housekeeping genes (HSKG) in resected tissue from TLE patients.** HSKG index in resected hippocampus (**a**) and amygdala (**b**) of each TLE patient (n=22). Indexes were calculated by integrating transcript levels of DMD, GAPDH and HPRT1 housekeeping genes. (**c**) Fold-difference of HSKG index between the amygdala and the hippocampus is given for each patient

**Fig. S3 The normalization techniques used in RT-qPCR can modify the results.** Comparison of the pro-inflammatory index (PI-I) values for each patient after unbiased normalization with the SmRNA (filled bars) or after normalization with housekeeping genes (dotted bars) in the hippocampus (**a**) and the amygdala (**b**)

**Fig. S4 Evolution of glial cell activation in the hippocampus after Pilo-SE.** Immunofluorescence detection was performed in the rat hippocampus using specific antibodies directed against ITGAM (CD11b) for microglia/macrophages (magenta) and GFAP for astrocytes (green). Nuclei were counterstained with DAPI. For GFAP, the rabbit polyclonal anti-GFAP antibody was used (AB5804; Chemicon). Different stages of epileptogenesis (SE-1D: 1-day post-SE; SE-9D: 9 days post-SE) or chronic epilepsy (SE-7W: 7 weeks post-SE) after Pilo-SE induced at weaning (P21) or at juvenile age (P42) are compared to their respective controls. Scale bar: 500 *µ*m

**Fig. S5 Increased expression of representative genes of the interleukin 1 system (IL1β, IL1R1, IL1RA) in the hippocampus of epileptic rats is dependent of the age at which SE is induced. a-b:** Immunohistochemical labeling of IL1β in the molecular layer of the hippocampus of Sprague-Dawley rats at the epileptic stage (7 weeks post-Pilo-SE; EPI 7W) after SE induced at juvenile age (**a2**, **a3**) or at weaning (**b2**, **b3**) and compared to their respective controls (CTRL 7W; **a1**, **b1**). IL1β protein is clearly detected in the hippocampus of epileptic rats subjected to SE at the juvenile stage. **c-d:** Transcript levels of IL1R1 (**c,** interleukin 1 receptor) and IL1RA (**d**, interleukin 1 receptor antagonist) were quantified once epilepsy was chronically installed, i.e. 7 weeks post-SE (EPI-W, n=8; EPI-J, n=8) compared to respective controls. Green asterisks indicate statistical significance between the two models (SE induced at weaning or juvenile stage), black asterisks indicate statistical significance between CTRL and SE. For IL1R1 in the EPI-J model, the statistical difference between CTRL and SE was p= 0.0723. Bonferroni *post-hoc* analysis following two-way ANOVA: ** *p*<0.01, *** *p*<0.001. Abbreviations: EPI-W, SE induced at weaning; EPI-J, SE in induced at juvenile stage. Scale bar: 50 *µ*m

**Fig. S6 RNAscope® ISH of IL1β-mRNA in resected hippocampus from TLE patients corroborates data obtained in the same hippocampus by RT-qPCR.** RNAscope® ISH of IL1β-mRNA (**a**, magenta) was detected together with ITGAM (CD11b)-mRNA (**b**, green) or GFAP-mRNA (**c**, green) in the resected hippocampus of TLE patients. Two patients are represented (P15 and P42) and the respective IL1β-cDNA copy numbers measured by RT-qPCR are provided. As shown by the white arrows, IL1β-mRNA appears to be located in cells bearing morphological features of glial cells. Scale bar: 50 *µ*m

**Fig. S7 RNAscope® ISH of IL1β, ITGAM and GFAP transcripts in the dentate gyrus of rats after Pilo-SE at 42 days.** Triple ISH of IL1β (**a**), ITGAM (CD11b) (**b**) and GFAP (**c**) transcripts using RNAscope® technology is depicted in the rat dentate gyrus. Nuclei were counterstained with DAPI. Different stages of epileptogenesis (SE-7H: 7 hours post-SE; SE-9D: 9 days post-SE) or chronic epilepsy (EPI-7W: 7 weeks post-SE) after Pilo-SE at juvenile age (P42) are compared to their respective controls (CTRL 7H/9D and CTRL 7W). Scale bar: 50 *µ*m

**Table S1 Primer sequences – Homo sapiens sapiens.** Abbreviations: DMD: Dystrophin; GAPDH: Glyceraldehyde 3-phosphate dehydrogenase; HPRT1: Hypoxanthine Phosphoribosyltransferase 1; IL1β: Interleukin 1 beta; IL6: Interleukin 6; TNF: Tumor necrosis factor; IFN*ɣ*: Interferon gamma; MCP1: Monocyte chemoattractant protein 1; MIP1⍺: Macrophage Inflammatory Protein alpha; IL4: Interleukin 4; IL10: Interleukin 10; IL13: Interleukin 13; GFAP: Glial fibrillary acidic protein; ITGAM: Integrin alpha M

**Table S2 Primer sequences – Rattus Norvegicus.** Abbreviations: As in Table S1; IL1R1: Interleukin 1 Receptor Type 1; IL1RA: Interleukin-1 receptor antagonist

**Table S3 Individual values of TLE patients for molecular and cellular markers of inflammation measured in the hippocampus.** Transcript level of pro-inflammatory cytokines (IL1β, IL6, TNF), chemokines (MCP1, MIP1⍺), anti-inflammatory cytokine IL10, and cell markers (GFAP, ITGAM) were measured in resected hippocampus of TLE patients (n=22). Individual values are expressed in percent of the mean value for each marker

**Table S4 Number of cDNA copies (mean ± SEM) in control rat hippocampus after reverse transcription of total RNA**

