## Supplementary material for "Low grade inflammation in the epileptic hippocampus contrasts with explosive inflammation occurring in the acute phase following *status epilepticus* in rats: translation to patients with epilepsy": N.Gasmi_Supplementary figures and tables

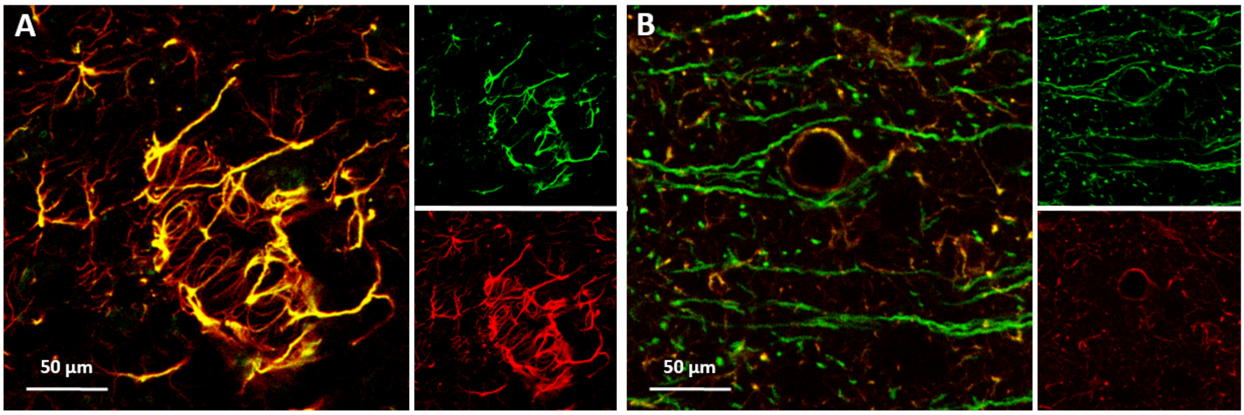


**Fig. S1 Non-overlapping of GFAP immunofluorescent labelling obtained with two different antibodies.** Double immunohistochemical labelling of GFAP in the hippocampus (**a**) and in the piriform cortex (**b**) of a rat 7 weeks after pilocarpine-induced SE. The rabbit polyclonal anti-GFAP antibody (AB5804; Chemicon) is visualized in green while the mouse monoclonal anti-GFAP antibody (G3893; Sigma-Aldrich) is visualized in red. Colocalization is displayed in yellow when red and green are superimposed. These observations suggest that GFAP epitopes recognized by the two antibodies are not accessible (or present) in the same manner within the same structure or between two different structures. Scale bar: 50 µm.

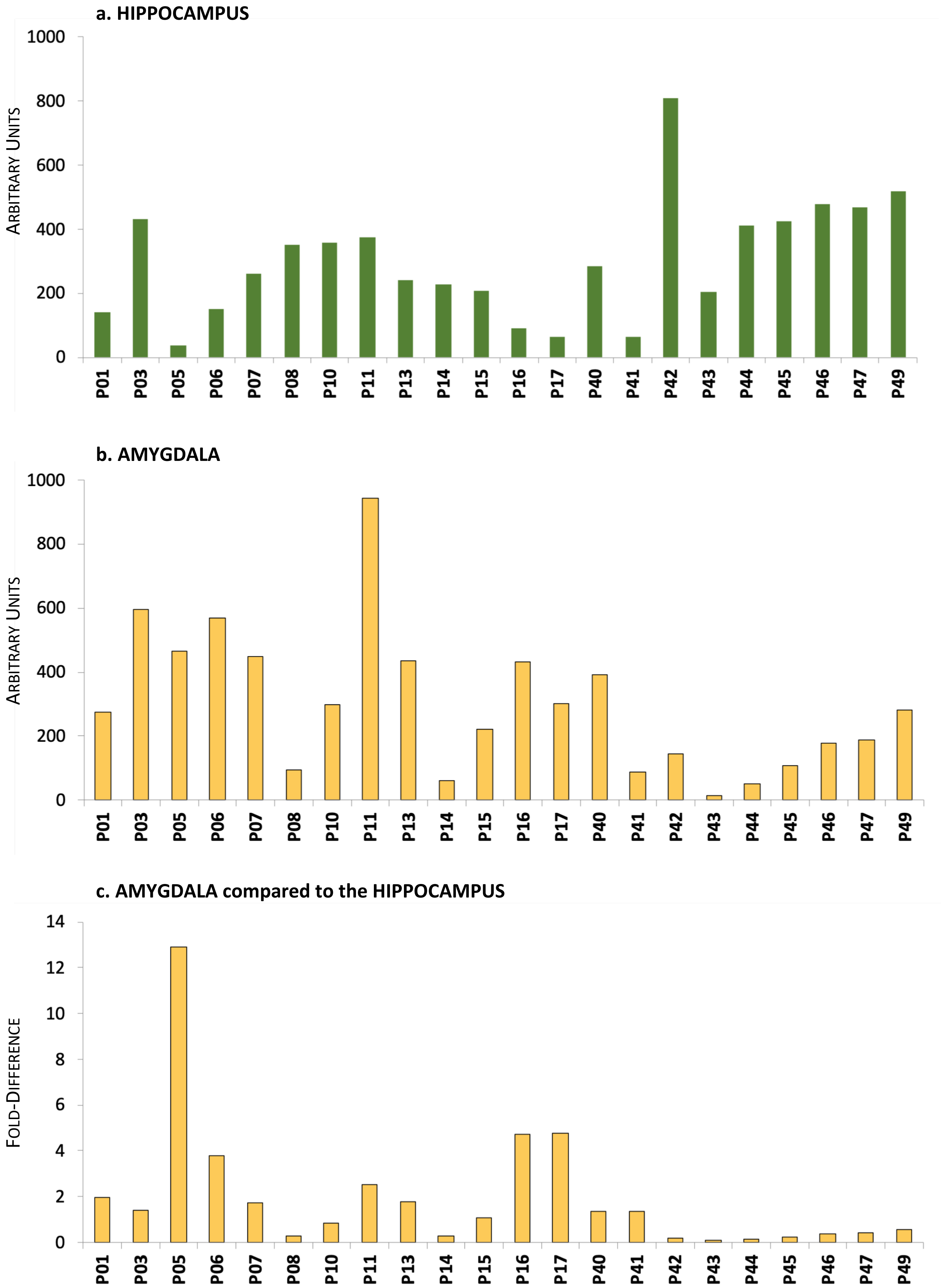


**Fig. S2 Index of housekeeping genes (HSKG) in resected tissue from TLE patients.** HSKG index in resected hippocampus (**a**) and amygdala (**b**) of each TLE patient (n=22). Indexes were calculated by integrating transcript levels of DMD, GAPDH and HPRT1 housekeeping genes. (**c**) Fold-difference of HSKG index between the amygdala and the hippocampus is given for each patient.

**a. HIPPOCAMPUS**

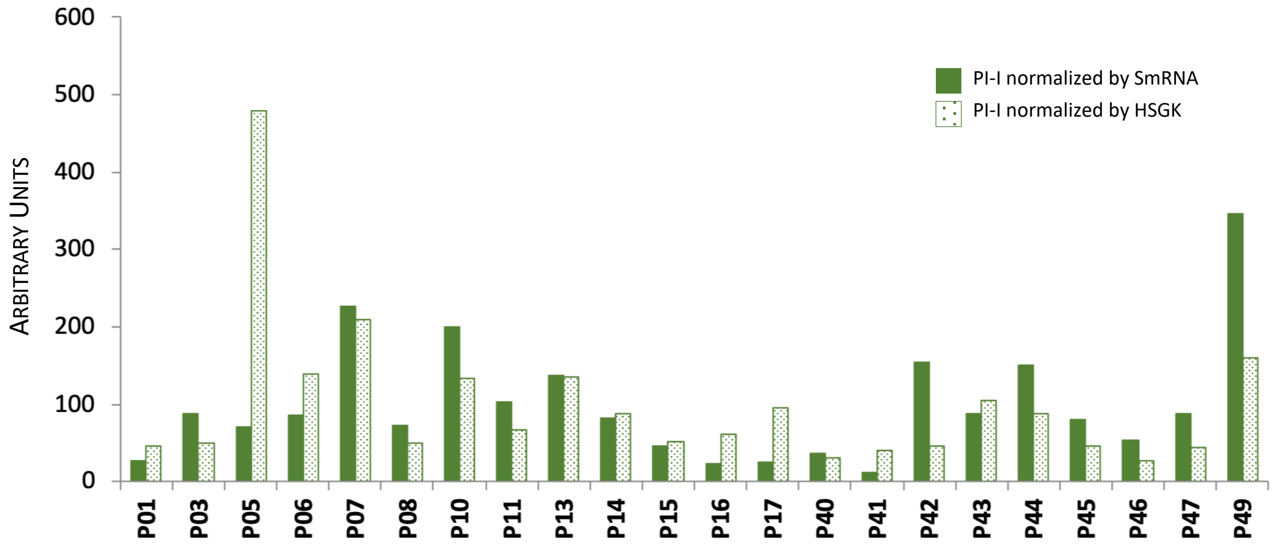


**b. AMYGDALA**

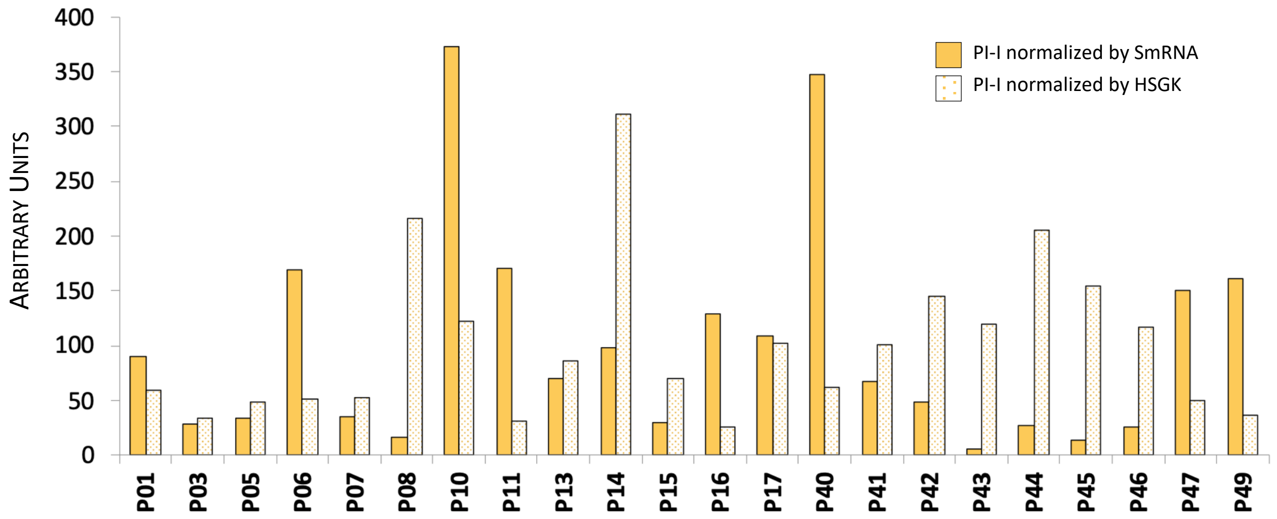


**Fig. S3 The normalization techniques used in RT-qPCR can modify the results.** Comparison of the pro-inflammatory index (PI-I) values for each patient after unbiased normalization with the SmRNA (filled bars) or after normalization with housekeeping genes (dotted bars) in the hippocampus (**a**) and the amygdala (**b**).

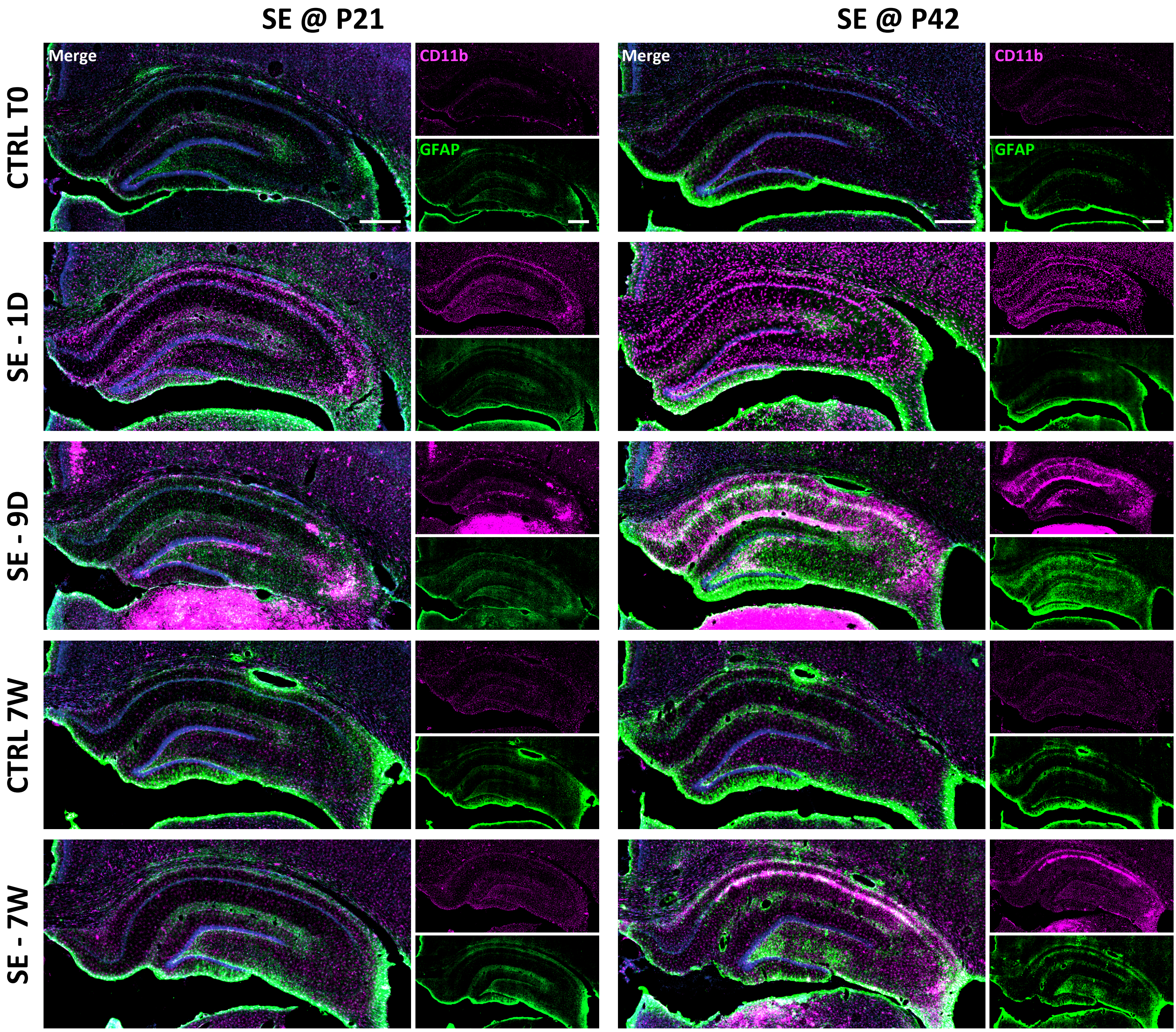


**Fig. S4 Evolution of glial cell activation in the hippocampus after Pilo-SE.** Immunofluorescence detection was performed in the rat hippocampus using specific antibodies directed against ITGAM (CD11b) for microglia/macrophages (magenta) and GFAP for astrocytes (green). Nuclei were counterstained with DAPI. For GFAP, the rabbit polyclonal anti-GFAP antibody was used (AB5804; Chemicon). Different stages of epileptogenesis (SE-1D: 1-day post-SE; SE-9D: 9 days post-SE) or chronic epilepsy (SE-7W: 7 weeks post-SE) after Pilo-SE induced at weaning (P21) or at juvenile age (P42) are compared to their respective controls. Scale bar: 500 $\mu$m.

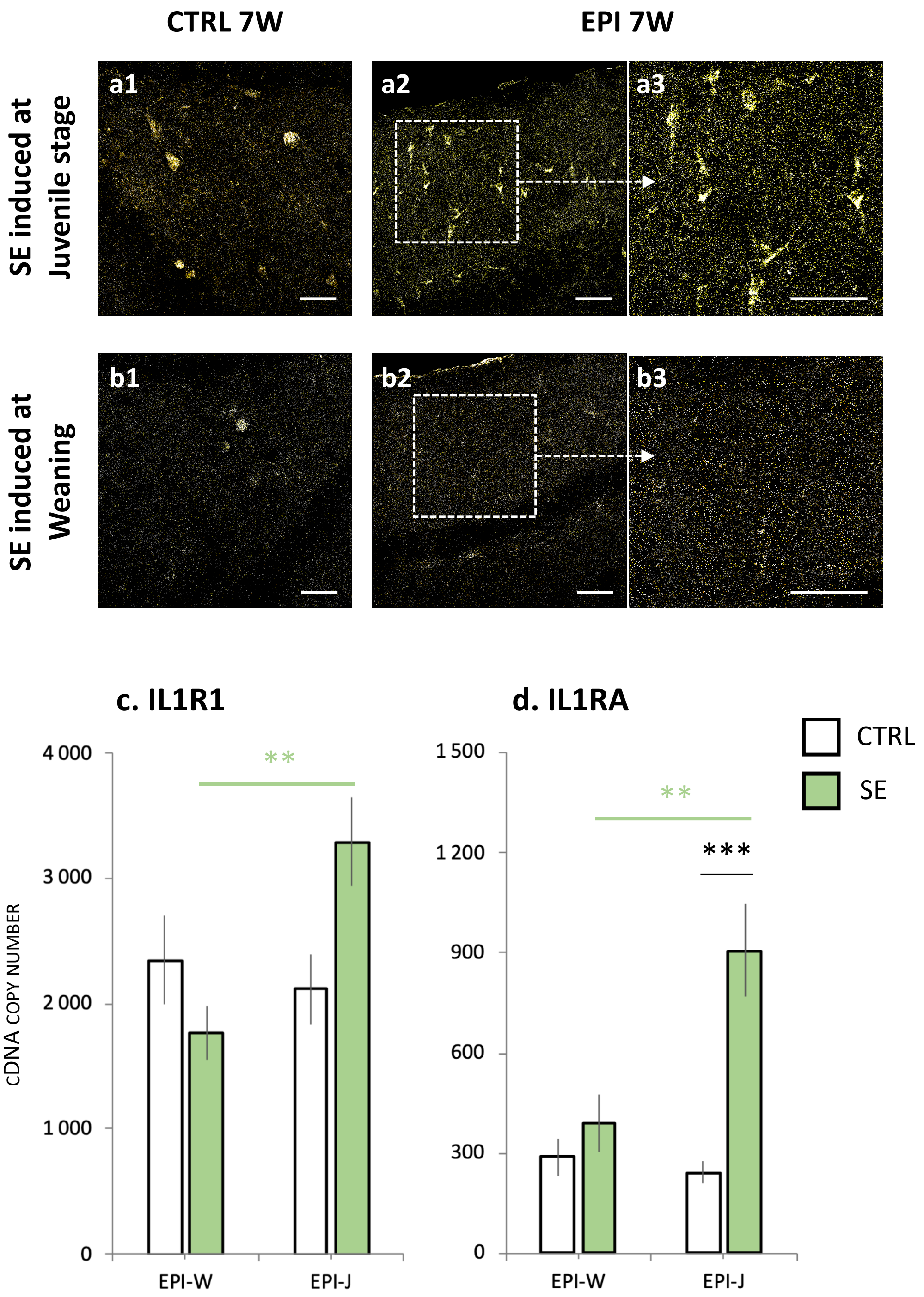


**Fig. S5 Increased expression of representative genes of the interleukin 1 system (IL1β, IL1R1, IL1RA) in the hippocampus of epileptic rats is dependent of the age at which SE is induced. a-b:** Immunohistochemical labeling of IL1β in the molecular layer of the hippocampus of Sprague-Dawley rats at the epileptic stage (7 weeks post-Pilo-SE; EPI 7W) after SE induced at juvenile age (**a2**, **a3**) or at weaning (**b2**, **b3**) and compared to their respective controls (CTRL 7W; **a1**, **b1**). IL1β protein is clearly detected in the hippocampus of epileptic rats subjected to SE at the juvenile stage. **c-d:** Transcript levels of IL1R1 (**c,** interleukin 1 receptor) and IL1RA (**d**, interleukin 1 receptor antagonist) were quantified once epilepsy was chronically installed, i.e. 7 weeks post-SE (EPI-W, n=8; EPI-J, n=8) compared to respective controls. Green asterisks indicate statistical significance between the two models (SE induced at weaning or juvenile stage), black asterisks indicate statistical significance between CTRL and SE. For IL1R1 in the EPI-J model, the statistical difference between CTRL and SE was p= 0.0723. Bonferroni *post-hoc* analysis following two-way ANOVA: ** $p<0.01$, *** $p<0.001$. Abbreviations: EPI-W, SE induced at weaning; EPI-J, SE in induced at juvenile stage. Scale bar: 50 $\mu$m.

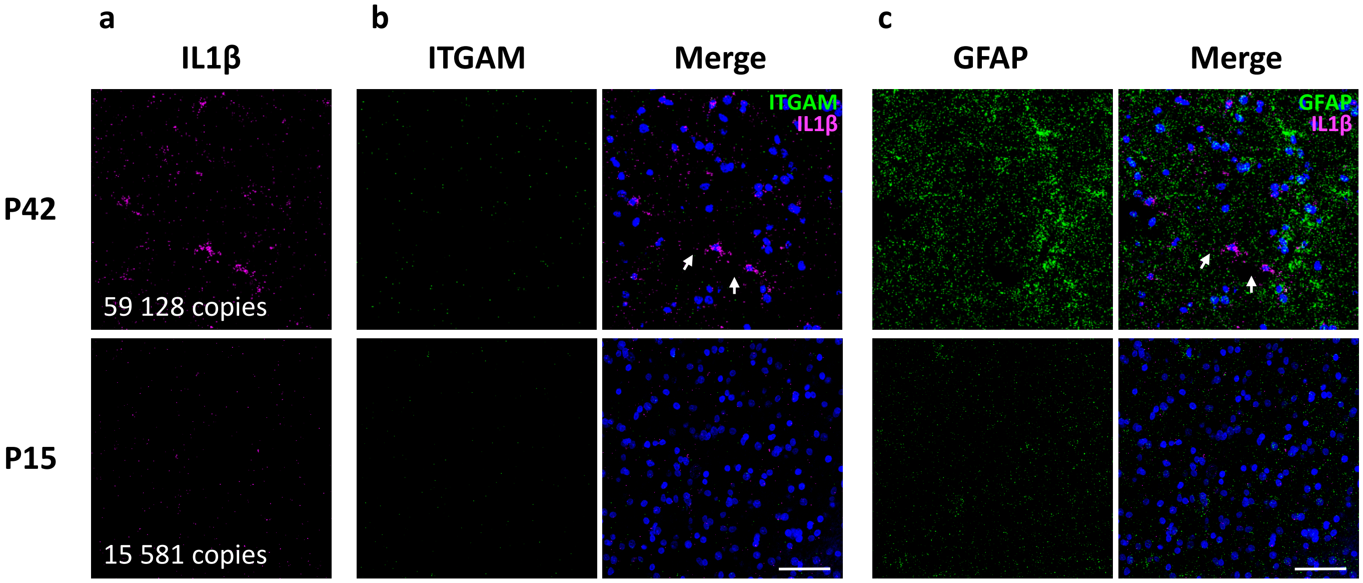


**Fig. S6 RNAscope® ISH of IL1β-mRNA in resected hippocampus from TLE patients corroborates data obtained in the same hippocampus by RT-qPCR.** RNAscope® ISH of IL1β-mRNA (**a**, magenta) was detected together with ITGAM (CD11b)-mRNA (**b**, green) or GFAP-mRNA (**c**, green) in the resected hippocampus of TLE patients. Two patients are represented (P15 and P42) and the respective IL1β-cDNA copy numbers measured by RT-qPCR are provided. As shown by the white arrows, IL1β-mRNA appears to be located in cells bearing morphological features of glial cells. Scale bar: 50 $\mu$m.

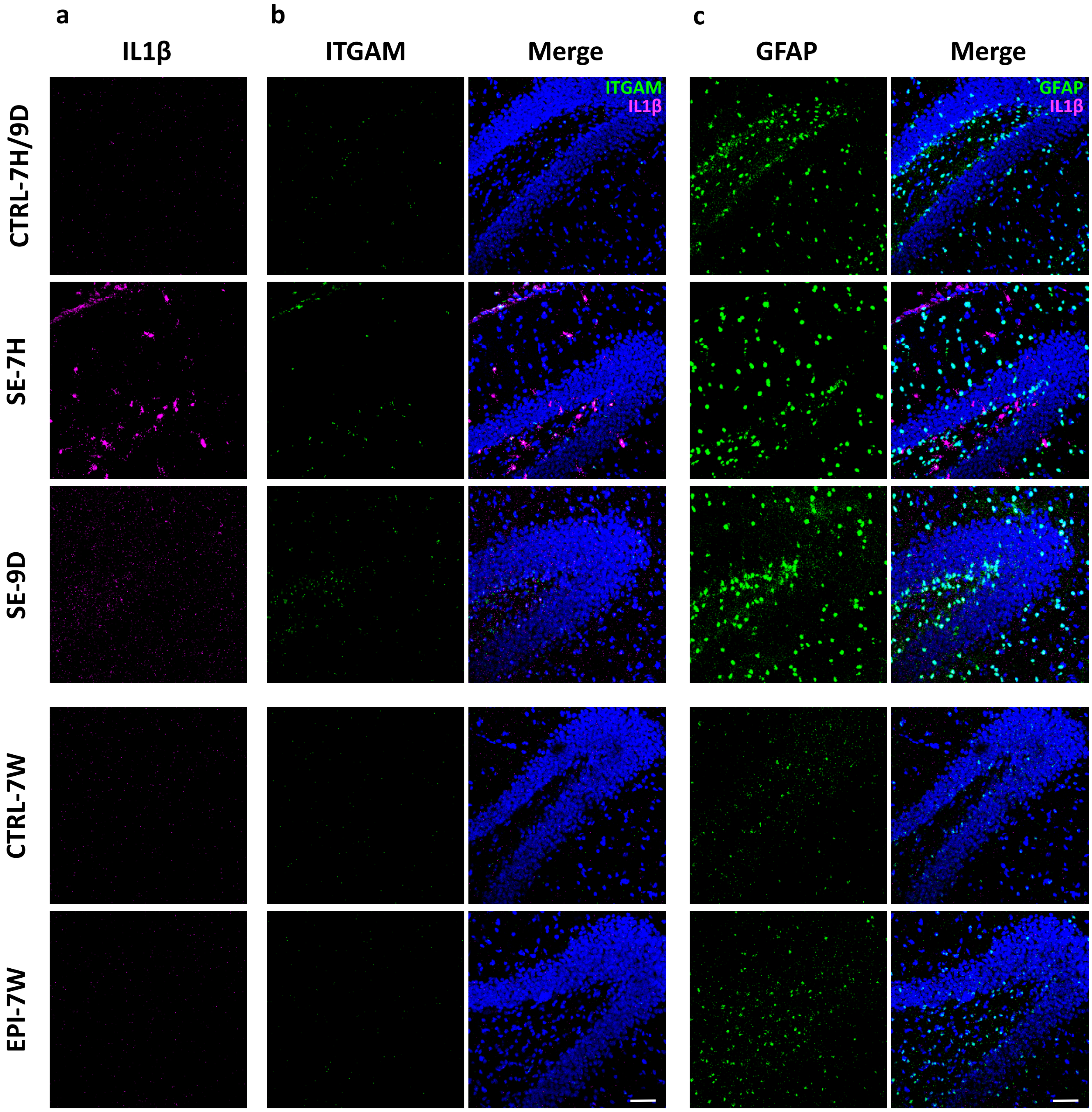


**Fig. S7 RNAscope® ISH of IL1β, ITGAM and GFAP transcripts in the dentate gyrus of rats after Pilo-SE at 42 days.** Triple ISH of IL1β (**a**), ITGAM (CD11b) (**b**) and GFAP (**c**) transcripts using RNAscope® technology is depicted in the rat dentate gyrus. Nuclei were counterstained with DAPI. Different stages of epileptogenesis (SE-7H: 7 hours post-SE; SE-9D: 9 days post-SE) or chronic epilepsy (EPI-7W: 7 weeks post-SE) after Pilo-SE at juvenile age (P42) are compared to their respective controls (CTRL 7H/9D and CTRL 7W). Scale bar: 50 $\mu$m.

| cDNA | Primer pairs – *Homo sapiens sapiens* | Product sizes (bp) | GenBank ID# |
|---|---|---|---|
| DMD | F-CCTCCACTCGTACCCACACT<br>R-TCCCAGCAAGTTGTTTGAGTC | 89 | NM_004015/<br>16/17/18.2 |
| GAPDH | F-AGCCACATCGCTCAGACAC<br>R-GCCCAATACGACCAAATCC | 66 | NM_002046.3 |
| HPRT1 | F-TGACCTTGATTTATTTTGCATACC<br>R-CGAGCAAGACGTTCAGTCCT | 102 | NM_000194.2 |
| IL1β | F-TACCTGTCCTGCGTGTTGAA<br>R-TCTTTGGGTAATTTTTGGGATCT | 76 | NM_000576.2 |
| IL6 | F-CAGGAGCCCAGCTATGAACT<br>R-AGCAGGCAACACCAGGAG | 85 | NM_000600.3 |
| TNF | F-CAGCCTCTTCTCCTTCCTGAT<br>R-GCCAGAGGGCTGATTAGAGA | 123 | NM_000594.2 |
| IFNγ | F-GGCATTTTGAAGAATTGGAAAG<br>R-TTTGGATGCTCTGGTCATCTT | 112 | NM_000619.2 |
| MCP1 | F-AGTCTCTGCCGCCCTTCT<br>R-GTGACTGGGGCATTGATTG | 93 | NM_002982.3 |
| MIP1α | F-TGCAACCAGTTCTCTGCATC<br>R-AATCTGCCGGGAGGTGTA | 75 | NM_002983.2 |
| IL4 | Qiagen®, Cat. #330001 PPH00565B | 93 | NM_000583.3 |
| IL10 | F-AGGACTTTAAGGGTTACCTGGGTTG<br>R-TTGATGTCTGGGTCTTGGTTCT | 103 | NM_000572.3 |
| IL13 | Qiagen®, Cat. #330001 PPH00688F | 78 | NM_002188.2 |
| GFAP | F-AGAGGGACAATCTGGCACA<br>R-CAGCCTCAGGTTGGTTTCAT | 71 | NM_002055.4 |
| ITGAM | F-GGCATCCGCAAAGTGGTA<br>R-GGATCTTAAAGGCATTCTTTCG | 70 | NM_000632.3 |

**Table S1 Primer sequences – *Homo sapiens sapiens.*** Abbreviations: DMD: Dystrophin ; GAPDH: Glyceraldehyde 3-phosphate dehydrogenase ; HPRT1: Hypoxanthine Phosphoribosyltransferase 1 ; IL1β: Interleukin 1 beta ; IL6: Interleukin 6; TNF: Tumor necrosis factor ; IFN$\gamma$: Interferon gamma ; MCP1: Monocyte chemoattractant protein 1 ; MIP1α: Macrophage Inflammatory Protein alpha ; IL4: Interleukin 4 ; IL10: Interleukin 10 ; IL13: Interleukin 13 ; GFAP: Glial fibrillary acidic protein ; ITGAM: Integrin alpha M.

| cDNA | Primer sequences – *Rattus norvegicus* | Product sizes (bp) | GenBank ID# |
|---|---|---|---|
| IL1β | F-TGTGATGAAAGACGGCACAC<br>R-CTTCTTCTTTGGGTATTGTTTGG | 70 | NM_031512.2 |
| IL1R1 | F- CACGGAATGAGACGATGGAAG<br>R- ACGAAGCAGATGAACGGATAG | 250 | NM_013123.3 |
| IL1RA | F-TCTGGAGATGACACCAAGCTC<br>R-GCCTCTAGTGTTGTGCAGAGG | 167 | NM_022194.1 |
| IL6 | F-CCCTTCAGGAACAGCTATGAA<br>R-ACAACATCAGTCCCAAGAAGG | 74 | NM_012589.1 |
| TNF | F-TGAACTTCGGGGTGATCG<br>R-GGGCTTGTCACTCGAGTTTT | 122 | NM_012675.3 |
| IFNγ | F-TTTTGCAGCTCTGCCTCAT<br>R-AGCATCCATGCTACTTGAGTTAAA | 107 | NM_138880.2 |
| MCP1 | F-CGGCTGGAGAACTACAAGAGA<br>R-TCTCTTGAGCTTGGTGACAAATA | 78 | NM_031530.1 |
| MIP1α | F-TCCACGAAAATTCATTGCTG<br>R-AGATCTGCCGGTTTCTCTTG | 92 | NM_013025.2 |
| IL4 | F-GTAGAGGTGTCAGCGGTCTG<br>R-TTCAGTGTTGTGAGCGTGGA | 70 | NM_201270.1 |
| IL10 | F-AGTGGAGCAGGTGAAGAATGA<br>R-TCATGGCCTTGTAGACACCTT | 62 | NM_012854.2 |
| IL13 | F-AGTCCTGGCTCTCGCTTG<br>R-GATGTGGATCTCCGCACTG | 63 | NM_053828.1 |
| GFAP | F-ACATCGAGATCGCCACCTAC<br>R-GGATCTGGAGGTTGGAGAAA | 90 | NM_017009.2 |
| ITGAM | F-ACTCTGATGCCTCCCTTGG<br>R-TCCTGGACACGTTGTTCTCA | 72 | NM_012711.1 |

**Table S2 Primer sequences – *Rattus Norvegicus.*** Abbreviations: As in Table S1; IL1R1: Interleukin 1 Receptor Type 1 ; IL1RA : Interleukin-1 receptor antagonist.

| Patient ID | IL1β | IL6 | TNF | MCP1 | MIP1α | IL10 | GFAP | ITGAM |
|---|---|---|---|---|---|---|---|---|
| **P01** | 24 | 26 | 14 | 31 | 38 | 11 | 19 | 32 |
| **P03** | 120 | 27 | 15 | 153 | 130 | 55 | 137 | 82 |
| **P05** | 18 | 25 | 220 | 61 | 34 | 137 | 17 | 6 |
| **P06** | 127 | 46 | 21 | 153 | 87 | 61 | 58 | 47 |
| **P07** | 80 | 559 | 93 | 120 | 280 | 77 | 124 | 95 |
| **P08** | 98 | 31 | 43 | 95 | 97 | 77 | 115 | 88 |
| **P10** | 407 | 98 | 29 | 189 | 273 | 287 | 146 | 109 |
| **P11** | 109 | 177 | 65 | 101 | 64 | 78 | 82 | 81 |
| **P13** | 154 | 179 | 32 | 202 | 117 | 208 | 122 | 54 |
| **P14** | 118 | 93 | 12 | 130 | 62 | 39 | 42 | 41 |
| **P15** | 55 | 26 | 41 | 66 | 40 | 44 | 91 | 47 |
| **P16** | 12 | 13 | 26 | 27 | 40 | 46 | 53 | 39 |
| **P17** | 30 | 0 | 0 | 64 | 32 | 2 | 32 | 7 |
| **P40** | 28 | 30 | 49 | 42 | 34 | 82 | 91 | 78 |
| **P41** | 16 | 8 | 0 | 19 | 14 | 0 | 28 | 11 |
| **P42** | 209 | 89 | 75 | 211 | 188 | 77 | 150 | 206 |
| **P43** | 123 | 31 | 38 | 95 | 156 | 9 | 151 | 135 |
| **P44** | 113 | 37 | 425 | 104 | 72 | 171 | 182 | 140 |
| **P45** | 62 | 33 | 207 | 56 | 49 | 260 | 153 | 247 |
| **P46** | 39 | 31 | 119 | 34 | 44 | 85 | 89 | 153 |
| **P47** | 58 | 43 | 126 | 41 | 170 | 92 | 133 | 260 |
| **P49** | 201 | 594 | 550 | 206 | 180 | 300 | 184 | 242 |

**Table S3 Individual values of TLE patients for molecular and cellular markers of inflammation measured in the hippocampus.** Transcript level of pro-inflammatory cytokines (IL1β, IL6, TNF), chemokines (MCP1, MIP1α), anti-inflammatory cytokine IL10, and cell markers (GFAP, ITGAM) were measured in resected hippocampus of TLE patients (n=22). Individual values are expressed in percent of the mean value for each marker.

| | P21 - Weaning | | | | P42 - Juvenile | | | | Pool P21/P42 | |
|---|---|---|---|---|---|---|---|---|---|---|
| | Epileptogenesis | | Epilepsy | | Epileptogenesis | | Epilepsy | | Epilepsy | |
| | 7H/1D/9D | | 7W | | 7H/1D/9D | | 7W | | 7W | |
| | Mean | SEM | Mean | SEM | Mean | SEM | Mean | SEM | Mean | SEM |
| **IL1β** | 106 | 13 | 247 | 28 | 199 | 16 | 195 | 28 | 221 | 21 |
| **IL6** | 0 | 0 | 0 | 0 | 15 | 5 | 0 | 0 | 0 | 0 |
| **TNF** | 12 | 5 | 37 | 2 | 32 | 5 | 36 | 6 | 37 | 3 |
| **IFNγ** | 0 | 0 | 14 | 4 | 0 | 0 | 11 | 3 | 13 | 2 |
| **MCP1** | 844 | 111 | 1 733 | 178 | 1 980 | 350 | 1 496 | 292 | 1 614 | 167 |
| **MIP1α** | 2 189 | 291 | 3 093 | 411 | 2 767 | 351 | 2 336 | 342 | 2 714 | 279 |
| **IL4** | 92 | 17 | 132 | 21 | 168 | 45 | 135 | 18 | 133 | 13 |
| **IL10** | 21 | 1 | 22 | 5 | 31 | 7 | 19 | 6 | 20 | 4 |
| **IL13** | 940 | 137 | 1 678 | 248 | 2 253 | 219 | 1 704 | 331 | 1 691 | 197 |
| **ITGAM** | 5 | 1 | 11 | 1 | 13 | 1 | 10 | 1 | 11 | 1 |
| **GFAP** | 228 476 | 8 908 | 372 334 | 9413 | 421 669 | 27720 | 380 725 | 14148 | 376 529 | 8199 |

**Table S4 Number of cDNA copies (mean ± SEM) in control rat hippocampus after reverse transcription of total RNA.**
